# Mapping the visual cortex with Zebra noise and wavelets

**DOI:** 10.1101/2025.07.19.665666

**Authors:** Sophie Skriabine, Maxwell Shinn, Samuel Picard, Kenneth D Harris, Matteo Carandini

**Affiliations:** University College London, London WC1E 6AE, United Kingdom

## Abstract

Studies of the early visual system often require characterizing the visual preferences of large populations of neurons. This task typically requires multiple stimuli such as sparse noise and drifting gratings, each of which probe only a limited set of visual features. Here we introduce a new dynamic stimulus with sharp-edged stripes called Zebra noise and a new analysis model based on wavelets, and show that in combination they are highly efficient for mapping multiple aspects of the visual preferences of thousands of neurons. We used two-photon calcium imaging to record the activity of neurons in the mouse visual cortex. Zebra noise elicited strong responses that were more repeatable than those evoked by traditional stimuli. The wavelet-based model captured the repeatable aspects of the resulting responses, providing measures of neuronal tuning for multiple stimulus features: position, orientation, size, spatial frequency, drift rate, and direction. The method proved efficient, requiring only 5 minutes of stimulus (repeated 3 times) to characterize the tuning of thousands of neurons across visual areas. In combination, the Zebra noise stimulus and the wavelet-based model provide a broadly applicable toolkit for the rapid characterization of visual representations, promising to accelerate future studies of visual function.

## Introduction

Neurons in the early visual cortex are selective for a wide range of visual attributes, and this selectivity is currently probed with multiple distinct synthetic stimuli. For instance, neurons in the primary visual cortex are selective for properties such as position, size, orientation, spatial frequency, speed and direction. Initial efforts to measure their selectivity were aimed at single neurons and focused on a few visual properties at a time, with stimuli such as bars (Hubel & Wiesel, 1959), squares (Jones & Palmer, 1987b), and gratings (Ringach et al., 1997, Movshon et al., 1978b). In later efforts, the emphasis shifted towards stimuli that can simultaneously characterize many aspects of selectivity in multiple neurons. These include contrast-modulated pink noise (Niell & Stryker, 2008), sums of spatiotemporal wavelets (Bonin et al., 2011), and structured noise stimuli such as “Monet” and “Trippy” (MICrONS Consortium et al., 2025).

An efficient stimulus for characterizing populations of visual neurons should elicit strong and repeatable responses while varying along multiple visual dimensions. The activity of neurons in the visual system is determined not only by their visual preferences but also by non-visual factors (Niell & Stryker, 2010, Erisken et al., 2014, Schröder et al., 2020). An ideal stimulus for mapping visual preferences would evoke responses that are stronger than this non-visual activity, and thus repeatable. Moreover, an ideal stimulus would minimize biases, to avoid compensatory adaptations that are typical of the visual system when stimuli are unbalanced (Benucci et al., 2009, Dhruv & Carandini, 2014). Fi-nally, an ideal stimulus should allow rapid mapping, so that recording sessions can be brief or mostly devoted to other purposes.

To meet these criteria, we designed a dynamic visual stimulus we call “Zebra noise”. The stimulus is made of stripes that vary randomly in orientation, width, eccentricity, direction, and speed. The stripes have high contrast and sharp edges. Because edges have power at multiple spatial frequencies, they can drive neurons with diverse spatial frequency preferences. Moreover, the sharp edges cause the spatial frequency components to be in phase, leveraging nonlinear effects that increase the responses of neurons in visual cortex (Felsen et al., 2005).

To analyze the responses of visual cortical neurons to Zebra noise, we designed a simple non-linear analysis based on Gabor wavelets (“WavEn”). The analysis is inspired by models of simple and complex cells in the primary visual cortex (Movshon et al., 1978a, Movshon et al., 1978b) and by their subsequent elaborations (Touryan et al., 2005, Vintch et al., 2015,de Vries et al., 2020). It is a basic approximation of known visual tuning properties, designed to characterize the visual preferences that are probed by Zebra noise.

In combination, the Zebra noise stimulus and the wavelet-based analysis proved highly effective in characterizing the preferences of neurons in the visual cortex. We used two-photon imaging to record the responses of thousands of neurons in the mouse visual cortex. Zebra noise elicited strong and repeatable responses both in the primary visual area and in higher vis-ual areas. These responses were well described by the wavelet-based model. Moreover, the stimulus and the analysis were highly efficient: even 5 minutes of Zebra noise, repeated 3 times, was sufficient to characterize the responses of thousands of neurons. These results suggest that Zebra noise and a wavelet-based analysis are useful tools to characterize the visual system.

## Results

Zebra noise contains black and white stripes of varying size, orientation, and velocity. A brief video is available at **[Supplementary Movie 1](https://drive.google.com/file/d/1CQpGF1nPTHnaDrirgJseSkrDc0_4T175/view?usp=drive_link)** and a longer one is [on YouTube](https://www.youtube.com/watch?v=-SyjgbNCP4Q)). To generate it, we start from 3-dimensional fractal Perlin noise (Perlin, 1985), which is computed rapidly on a pixel-by-pixel basis (**Figure 1a**). We then apply a “comb threshold” to the gray values, discretizing it into bins of equal size, and setting the pixels to white if their value falls into an even bin, and to black otherwise (**Figure 1b**). Decreasing the number of teeth in the comb (increasing the bin size) creates patterns with thicker stripes (**Figure 1b,c**). In addition to the bin size, Zebra noise has two other tunable parameters: the fractal exponent of the Perlin noise, which controls the jaggedness of the contours, and the zoom magnitude, which controls the overall scale (Supplementary Figure 1). The movement of the stripes creates a constantly changing motion field (**Figure 1d**). The resulting movie is largely uncorrelated over space (**Figure 1e**). It contains power in multiple spatial frequencies and is approximately uniform across orientations (**Figure 1f,g**).

**Figure 1.**
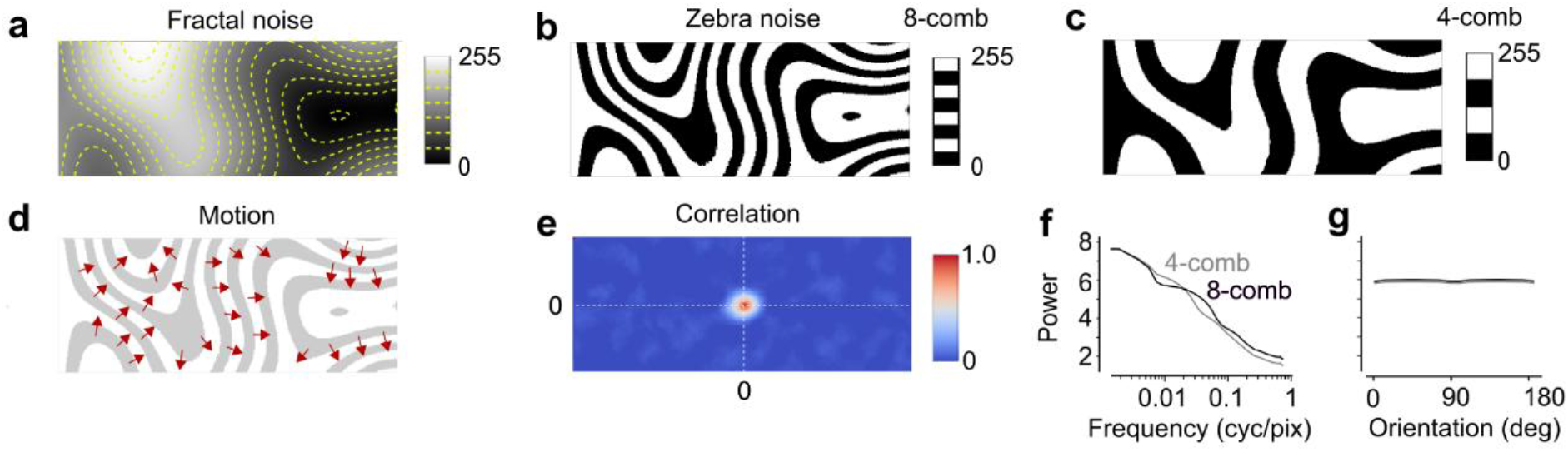
The Zebra noise stimulus. **a**: A frame of fractal Perlin noise, with contours at 8 gray levels (yellow curves). **b**: The resulting frame of Zebra noise, obtained by applying an 8-tooth comb threshold to the intensity scale. **c**: A coarser version of the Zebra noise obtained by applying a 4-tooth comb threshold to the same frame of 1/f noise. **d**: Vector field showing the movement of the edges in the Zebra noise in b. For clarity, the spatial pattern is shown at reduced contrast. **e**: Spatial autocorrelation of the Zebra noise movie. **f**: Spatial frequency content of the Zebra noise stimuli in b,c. **g**: Orientation content of the two Zebra noise stimuli. The quantities in e-g were computed over 1,500 stimulus frames.

We tested the Zebra noise stimulus in mouse visual cortex and found that it elicited strong and repeatable responses. We performed recordings with a two-photon mesoscope (Sofroniew et al., 2016) in the visual cortex of transgenic mice expressing the calcium indicator GCaMP6s in excitatory neurons (Wekselblatt et al., 2016) (**Figure 2a-c**). When we presented Zebra noise (7 min, repeated 3 times), the neurons gave strong responses (e.g. **Figure 2d**). To assess response repeatability, we measured the average correlation across repeats. The responses of many neurons were highly repeatable, both in primary visual cortex and in higher areas (**Figure 2e**).

**Figure 2.**
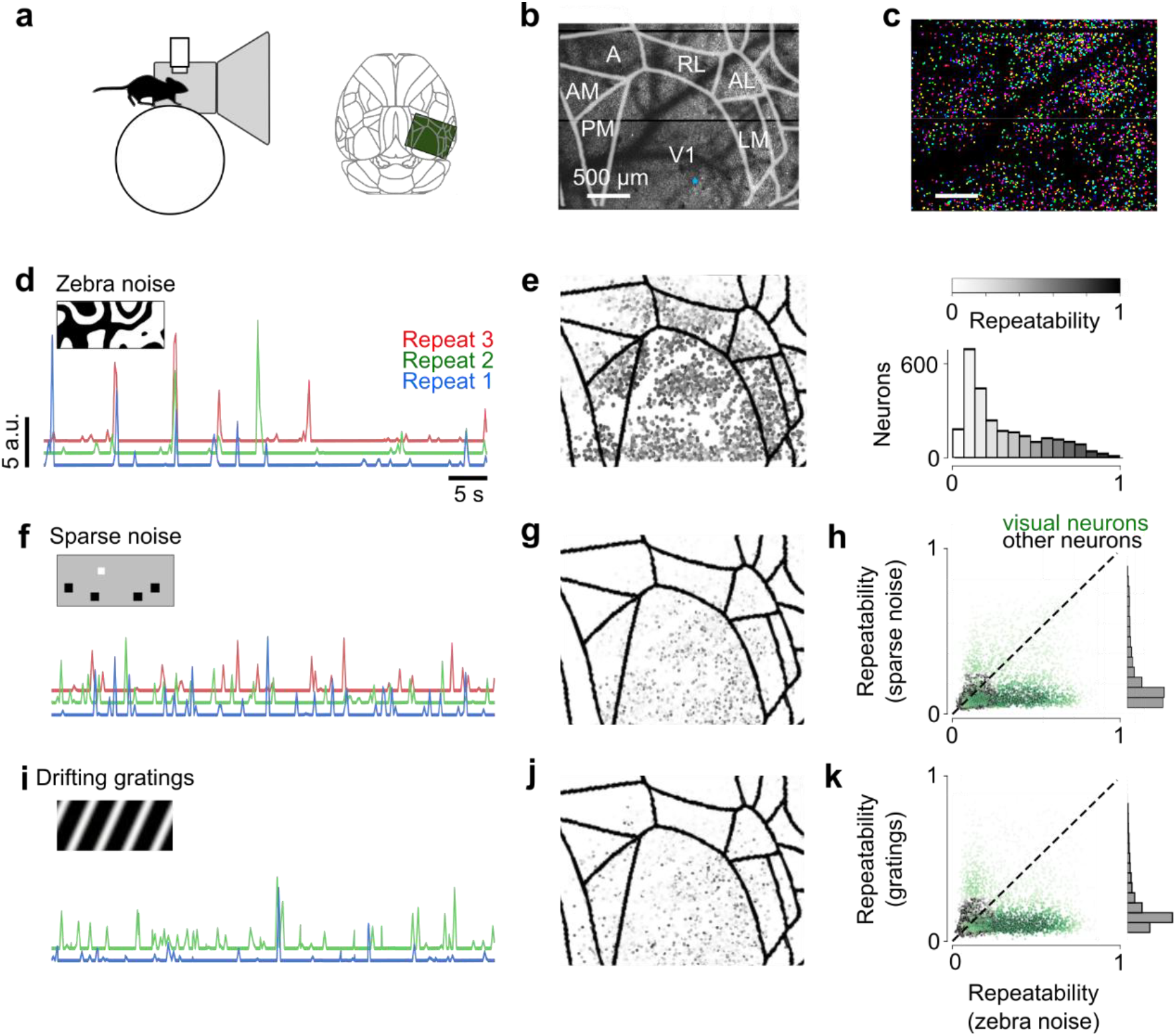
Zebra noise elicits repeatable responses in the mouse visual cortex. **a**: Head-fixed mice were free to run on a wheel and viewed images on two screens, in front and on the left, while a 2-photon microscope imaged the right visual cortex (*green rectangle*). **b**, Mean fluorescence image from the 2-photon microscope. Curves outline primary visual cortex (V1) and higher visual areas (HVA), as defined by the Allen Mouse Brain Common Coordinate Framework (Wang et al., 2020). **c:** Neurons detected in the field of view (with random colors). **d**: Responses of an example neuron to three repeats of Zebra noise. *Scale bar* shows 5 arbitrary units of deconvolved firing rate. **e**: Repeatability across repeats of the responses of the neurons in the field of view for Zebra noise. **f**: Responses of the same neuron as d to sparse noise stimuli. **g**, Repeatability of neuronal responses for sparse noise. **h**: Comparison of repeatability of responses to sparse noise compared to Zebra noise in neurons that gave repeatable visual responses (*green*) and remaining neurons (*black*). **i-k**: Same as f-h, for drifting grating stimuli.

These responses were more repeatable than those elicited by visual stimuli that are commonly used to map selectivity for stimulus position (sparse noise) or for stimulus orientation and direction (drifting gratings). In the same imaging session, we presented sparse noise made of black and white flashed squares, a visual stimulus commonly used to map preferences for stimulus position (10 min, repeated 3 times). Responses elicited by sparse noise were generally less repeatable (e.g. **Figure 2f**). Among the neurons that showed repeatable visual responses (n = 2,026 / 3,907 neurons with repeatability > 0.2 with any stimulus), 70% had higher repeatability in response to Zebra noise than in response to sparse noise (**Figure 2g-h**). We also showed drifting gratings (18 directions in 5 min, repeated twice), the visual stimulus typically used to measure preferences for stimulus orientation and direction. Once again, we observed responses that were substantially less repeatable throughout the visual cortex and especially in higher visual areas (**Figure 2i-j**). Among the neurons with repeatable visual responses, 75% had higher repeatability in response to Zebra noise than to drifting gratings (**Figure 2k**). Therefore, Zebra noise elicited more repeatable visual responses than sparse noise and drifting gratings.

Zebra noise also elicited more repeatable responses than “Trippy”, a similar dynamic stimulus that appears blurrier. Like Zebra noise, Trippy has irregular and dynamic bands, but it lacks sharp edges (MICrONS Consortium et al., 2025; Yatsenko et al., 2018). We compared the responses of V1 neurons in to Zebra noise and to Trippy and found the latter to be less repeatable (Supplementary Figure 2), suggesting that the sharp edges of Zebra noise enhance the repeatability of the neural responses.

Zebra noise triggered a repeatable response also in neurons that fired rarely. Some neurons fired only a few times during 7 minutes of Zebra noise and thus had a distribution of firing rates that was highly skewed. The responses of these neurons were highly repeatable (repeatability > 0.5, Supplementary Figure 3). By contrast, the same neurons gave much less repeatable responses to drifting gratings and sparse noise, with a correlation across trials typically <0.2. This suggests that by sampling a rich parametric space, Zebra noise can reliably drive neurons that are highly selective and thus typically silent.

To analyze responses of neurons in primary visual cortex to Zebra noise, we started by filtering the stimulus with Gabor wavelets and pooling the results across phases. Gabor wavelets are a reasonable description of the receptive fields of V1 simple cells (Jones & Palmer, 1987a), and are thus a robust starting point for more elaborate models of cortical responses (de Vries et al., 2020; Movshon et al., 1978b, 1978a; Touryan et al., 2005; Vintch et al., 2015; Yoshida & Ohki, 2020). They are parametrized by their position x and y, orientation θ, size s, and spatial frequency f (**Figure 3a,b**). In addition, their phase φ can range from 0 to 2π. Filtering a frame of the Zebra noise stimulus with a Gabor wavelet highlights its edges along a given orientation (**Figure 3c-d**). We then combine the output of two wavelets in cosine phase (φ =0) and sine phase (φ = π/2) into a complex number and take its amplitude *A*, obtaining an image that highlights the edges regardless of their polarity (**Figure 3e**).

**Figure 3.**
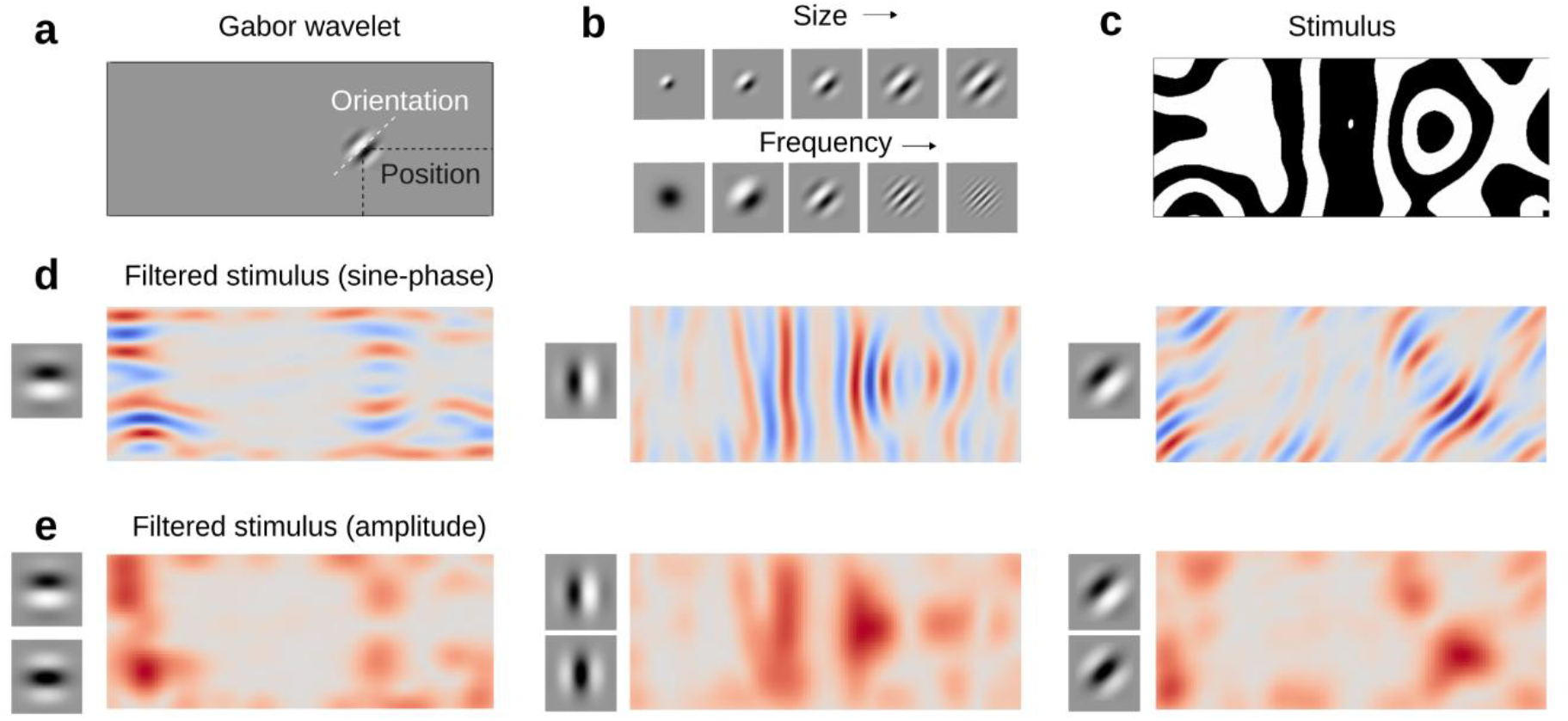
Wavelet transform of a Zebra noise video. **a**. Each Gabor wavelet has a position, orientation, size, and frequency. One example wavelet is shown. **b**. Examples of Gabor wavelets differing in size and frequency. **c**. A frame of Zebra noise. **d**. The result of filtering the stimulus frame with a sine-phase Gabor wavelet of intermediate scale and frequency, at three orientations. Red indicates positive values, blue negative values. **e**. The amplitude of the filtered image, obtained by combining the output of the sine-phase and cosine-phase Gabor wavelets.

For each neuron, we next selected the wavelet whose phase-pooled output best correlates with the neuron’s responses. Having obtained for each wavelet a filtered amplitude *A*(*t*) of the stimulus as a function of time *t*, we find for each neuron the wavelet whose *A*(*t*) best correlates with the activity of the neuron. The parameters of this best-fitting wavelet estimate the neuron’s preferred azimuth x, elevation y, and orientation θ (**Figure 4a**) as well as its preferred size s and spatial frequency f. For the example neuron (**Figure 4b**), the correlation between the neuron’s response and the amplitude *A*(*t*) of the best-fitting wavelet is 0.26 (**Figure 4c**).

**Figure 4.**
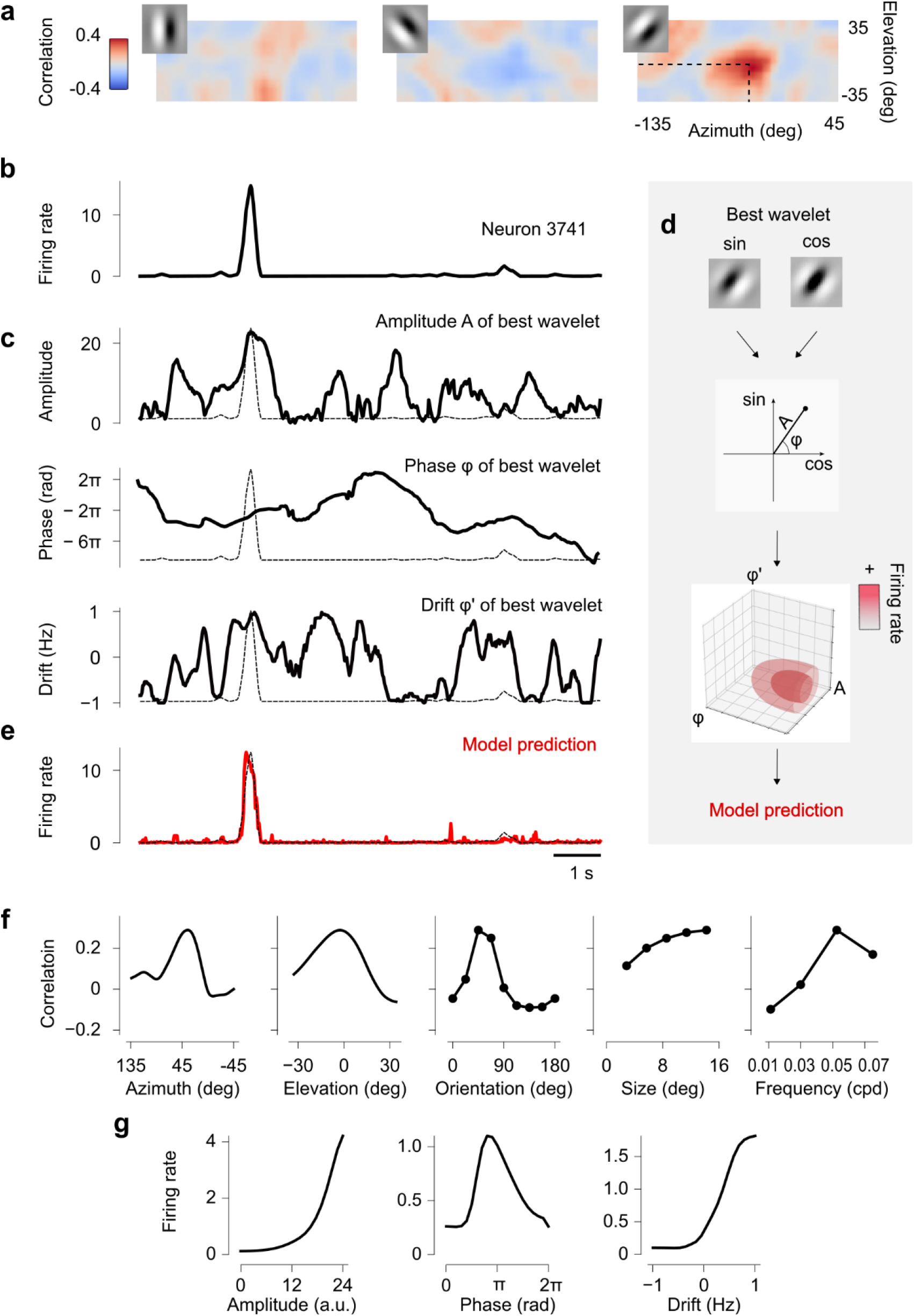
Fitting the wavelet-based model and obtaining tuning curves. **a**: Correlation between the responses of an example neuron and the wavelet transform of the stimulus, as a function of wavelet position, for 3 example wavelet orientations (*insets*). Correlation is strongest for the 45 deg orientation at a particular position (*dashed lines*). **b**: Firing rate of the example neuron during an interval of Zebra noise. **c**: Amplitude, phase, and phase derivative of the response of the best-fitting Gabor wavelet as a function of time, compared to the neuron’s firing rate (*dashed*). **d**: Cartoon of the model. The image is filtered with two versions of the best-fitting wavelet, in sine and cosine phase. The two outputs are then projected into a three-dimensional space with coordinates amplitude *A*, phase *φ*, and drift rate *φ*′ (the derivative of phase), providing a predicted response for each combination of these 3 values. **e**: Output of the model for the example neuron, compared to the measured firing rate (*dashed curve*). **f**: Tuning of the example neuron for azimuth, elevation, orientation, size (radius), and frequency. **g**: Tuning of the example neuron for wavelet amplitude, phase, and drift rate, showing that this neuron was highly selective for stimulus direction.

Finally, we refined the predictions by identifying for each neuron any nonlinear dependence on amplitude, phase, and phase derivative (drift rate) (**Figure 4d**). Having identified a best-fitting wavelet for each neuron, we go back to the complex-valued output of the sine-phase and cosine-phase wavelets and consider not only the amplitude *A*(*t*), but also the phase *φ*(*t*) and its time derivative *φ*′(*t*). We term the latter the drift rate; it represents both the direction and the speed of movement. At each moment *t*, these three quantities describe a point in a three-dimensional space, and the neuron’s response provides a value *R*(*A, φ, φ*′) in that point (**Figure 4d** and Supplementary Figure 4). As Zebra noise samples much of this three-dimensional space, we can obtain robust estimates for the value *R* for the entire space (Methods). This allows the model to predict responses to stimuli that were not strictly part of the training set. The resulting prediction is quite close to the actual activity of the neuron (**Figure 4e**). For instance, in the example neuron the correlation between responses and model prediction is 0.68 (**Figure 4e**).

This approach yields approximate tuning curves for multiple stimulus parameters. In the first stage in the model (**Figure 4a**), we estimate preferred azimuth x, elevation y, orientation θ, size s, and spatial frequency f by finding the wavelet that provides the highest correlation. We can then vary one parameter at a time and obtain a “tuning curve” showing how the correlation decreases when the parameter deviates from optimal (**Figure 4f**). In the second stage of the model (**Figure 4d**) we estimate a joint function *R*(*A, φ, φ*′) indicating how the response varies with amplitude *A*, phase *φ*, and drift rate *φ*′. To estimate a neuron’s selectivity for these attributes we take averages over two of the three parameters to obtain a (generally nonlinear) input/output function *F*(*A*); a tuning for spatial phase *P*(*φ*) and a tuning for direction and drift rate *S*(*φ*′) (**Figure 4g**). This analysis provides insight into how the stimulus attributes contribute to each neuron’s response.

Similar results were obtained across visually responsive neurons with different types of selectivity (Supplementary Figure 5). In ~80% of neurons, the tuning curves displayed a single well-defined peak, enabling straightforward estimation of position x and y, orientation θ, scale s, and spatial frequency f (similar to **Figure 4a**). However, in the remaining ~20%, the curves exhibited multiple peaks in x or y, complicating receptive field localization. To address this ambiguity, we introduced an optional correction step, where the user can opt to rely on the smooth organization of retinotopic maps. This allows the preferred position of the ambiguous neurons to be determined from the tuning preferences of adjacent neurons. However, this assumption may obscure the local scatter of preferred positions where this scatter is present (Bonin et al., 2011).

Having characterized the tuning of individual neurons with Zebra noise, we can now create maps of those tuning properties. The maps of retinotopy displayed the expected orthogonal representation of visual field azimuth and elevation (Dräger, 1975) with some degree of scatter (Bonin et al., 2011) (**Figure 5a-b**). These maps were similar to those obtained with traditional stimuli such as sparse noise and drifting gratings (Supplementary Figure 6). The maps of preferred spatial frequency and size, in turn, displayed slow variations across space (**Figure 5c-d**), with V1 neurons showing, on average, smaller receptive fields. The map of preferred orientation exhibited the typical salt-and-pepper organization (Ohki & Reid, 2007) (**Figure 5e**). As is customary, from the retinotopy maps, we extracted the sign map, which encodes the sign of retinotopic gradients (**Figure 5f**). We used the sign map to delineate visual areas (Zhuang et al., 2017) (curves in **Figure 5a-g**).

**Figure 5.**
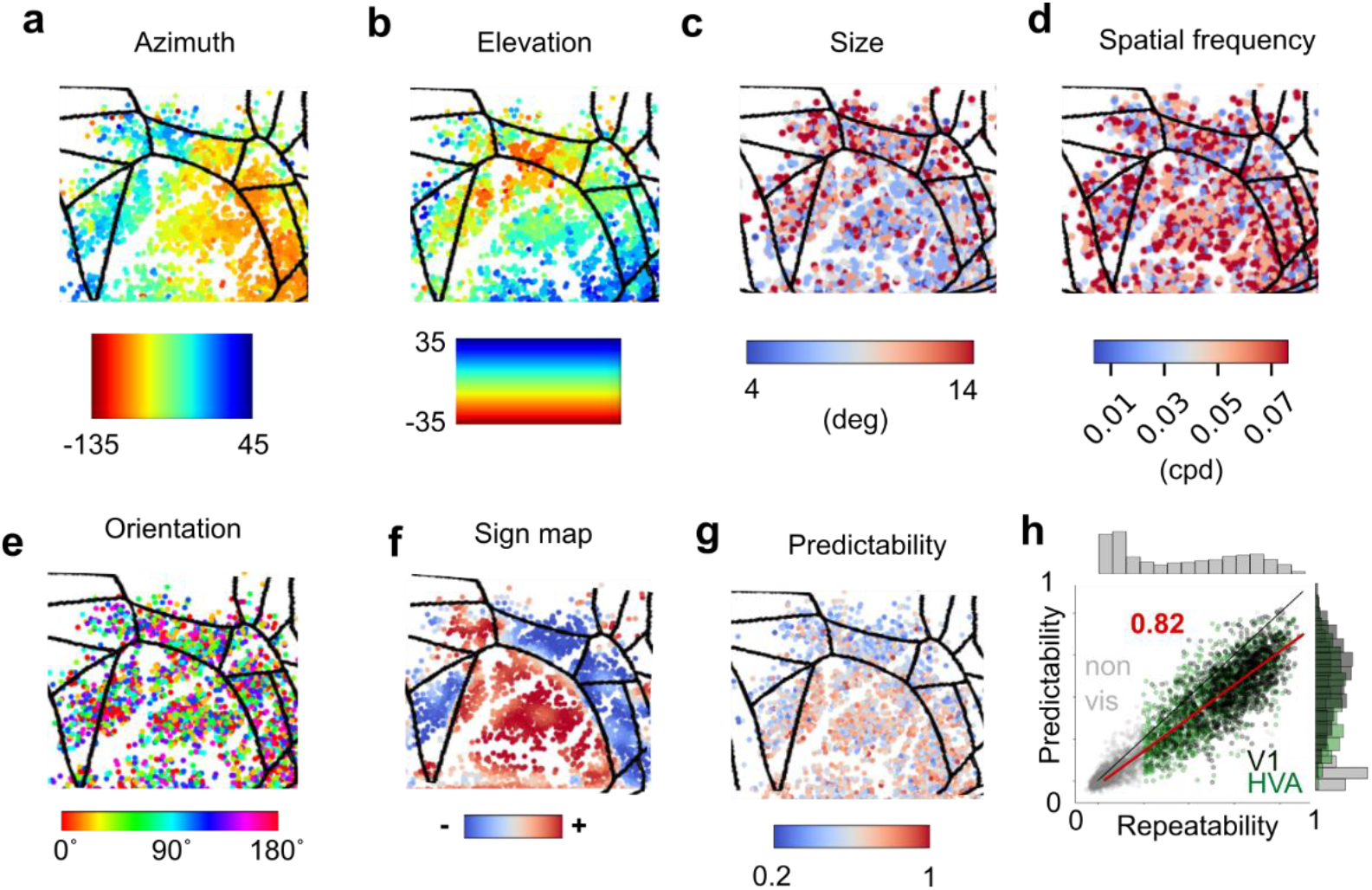
Mapping of neural tuning parameters and model predictive quality. **a:** Map of preferred azimuth in the visual cortex of an example mouse. Each dot is a neuron. Its color is the preferred azimuth in visual degrees. The different visual areas are delineated by the black contours. **b-e:** Same for preferred elevation (b), size (radius) (c), spatial frequency (d), and orientation (e). **f:** Sign map, showing areas where a clockwise rotation in cortex marks a clockwise (*red*) or counter-clockwise (*blue*) rotation in the visual field. **g:** Map of the predictability of the neuron. The color shows the cross-validated correlation between the model prediction and the neuron’s firing rate. **h:** Relation between predictability by the model and repeatability of a neuron’s firing rate across repeats of the same stimulus. Each dot is a neuron, colored according to whether it had high repeatability and was in V1 (*black)* or higher visual areas (*green*) or it had low repeatability (<0.2) (*gray*). The marginal distribution for repeatability and predictability are shown on the sides. The red line represents the linear fit. Its slope (0.82) indicates that the model captures the aspects of the responses that are repeatable across trials.

The model performed well both in primary visual cortex and in higher visual areas. To validate the model, we measured the correlation (cross-validated) between the model’s predictions and neural activity (predictability). Predictability was generally higher for V1 neurons, with an average of 0.44, and slightly lower for higher visual areas (HVA) neurons, with an average of 0.36 (**Figure 5g**). However, this difference could be due to the higher repeatability (inter-trial correlation) of V1 responses, which we have already documented (**Figure 2e**). To assess this possibility, we computed a prediction score, defined as the ratio between predictability and repeatability. This score estimates the ability of the model to explain repeatable aspects of the responses. The model achieved a prediction score of 0.82 (**Figure 5h**) over all visually responsive neurons, indicating that it captured a large fraction of the visual properties of the neurons. This score (0.82) was on average the same for neurons in V1 and in HVAs, suggesting that the difference in predictability between V1 and HVA neurons is entirely explained by the difference in repeatability.

The Zebra noise stimulus proved highly efficient, such that 5 minutes were sufficient to provide a reasonable characterization of the visual properties of the neurons. Indeed, 5 minutes of Zebra noise (repeated 3 times) were sufficient to characterize retinotopic preferences, with no apparent improvement from adding an additional minute (**Figure 6a,b**). Similarly, 5 minutes were sufficient to characterize orientation preference, with minimal improvement by adding an additional minute (**Figure 6c,d**). However, for more complete characterizations of visual responses, it may be useful to record for longer. Indeed, the ability to predict responses to unseen stimuli kept on improving up to 6 minutes, with no sign of deceleration (Supplementary Figure 7).

**Figure 6.**
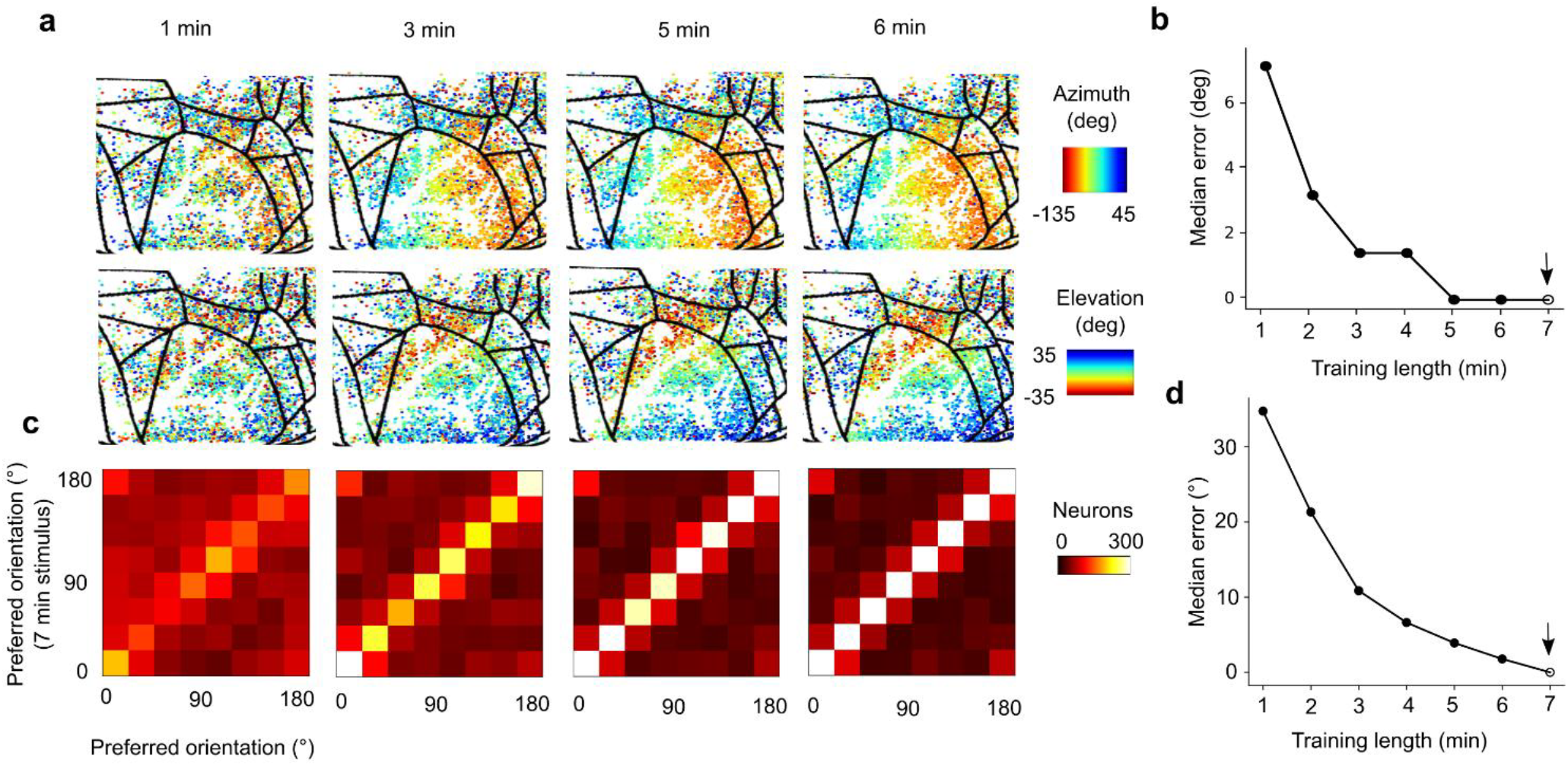
Effect of stimulus duration. **a:** Maps of preferred azimuth and elevation in the example mouse in Figure 5, for different durations of Zebra noise stimulus as indicated. The results for the full 7 minutes of stimulus are shown in Figure 5a,b. **b:** Median error in estimated retinotopic position, relative to the full 7-minute stimulus (*arrow*). **c**: Comparison of orientation preference obtained with the full 7-minute stimulus (*ordinate*) vs. shorter stimulus (*abscissa*), for different stimulus durations as indicated. The map obtained with the full 7-minute stimulus is shown in Figure 5e. **d**: Median error in estimated orientation preference, relative to the full 7-minute stimulus (*arrow*).

Similar results were obtained in other mice and across calcium indicators and expression strategies. The experiments described up to here were performed in a mouse from a transgenic line (TetO) expressing the calcium indicator GCaMP6s in excitatory neurons (Wekselblatt et al., 2016). To validate the approach in other preparations, we replicated our experiments in two additional mice: one from the same TetO line and one from a mouse line (RiboL) characterized by a soma localized expression of a ribosome-tethered GCaMP8m. Thanks to the soma-localized expression, this mouse line has minimal neuropil contamination. Moreover, GCaMP8m has faster calcium dynamics than GCaMP6s (Zhang et al., 2023). Zebra noise and our analysis method provided an efficient characterization of the visual cortex in both mice, with the model achieving a prediction score of 0.72 in the RiboL mouse and 0.67 in the TetO mouse (Supplementary Figure 8). These results indicate that in combination, the Zebra noise stimulus and our analysis method are effective for multiple calcium indicators and expression strategies.

## Discussion

Characterizing the tuning of large neuronal populations is a central challenge in visual neuroscience, which is hampered by stimuli that are inefficient or probe a limited feature space. Here, we have introduced a solution: a combination of a novel dynamic stimulus, Zebra noise, and an interpretable wavelet-based model. We demonstrated that the sharp, high-contrast edges of Zebra noise drive highly repeatable and robust responses across visual cortex, enabling the efficient mapping of multiple feature preferences in thousands of neurons from only a few minutes of data.

The success of Zebra noise is likely due not only to its high-contrast moving patterns of appropriate spatial and temporal scales, but also its sharp edges, which exert a powerful influence on neurons in the visual cortex (Felsen et al., 2005). The benefit of sharp edges appeared to be confirmed by our observation that the ‘Trippy’ stimulus (MICrONS Consortium et al., 2025), which has similar dynamic bands but lacks sharp edges, elicited responses that were less repeatable. Furthermore, by systematically sampling a rich parametric space (orientation, speed, etc.), Zebra noise reliably drove even neurons that were highly selective and fired sparsely, which are often missed by less comprehensive stimuli.

Analyzing responses to a rich, dynamic stimulus like Zebra noise requires a quantitative model. To this end, we developed a nonlinear wavelet-based model that occupies a middle ground between simple filters and complex deep neural networks. While advanced neural network models may achieve higher predictive accuracy (e.g., Cadena et al., 2019; Du et al., 2025; Tong et al., 2023; Wang et al., 2025; Willeke et al., 2023; Yamins et al., 2014), our approach prioritizes computational efficiency and interpretability. We showed that despite its simplicity, this model successfully provided good estimates of neuronal visual preferences across eight dimensions, defined by the azimuth and elevation of the stimulus position, and by stimulus orientation, size, spatial frequency, phase, magnitude, and drift.

Remarkably, the model performed equally well in mouse primary visual cortex (V1) and in the surrounding higher visual areas (HVA). Indeed, while predictability was slightly lower in HVA than in V1, this decrease in predictability was fully explained by a decrease in repeatability. Indeed, the ratio of predictability to repeatability (the prediction score) was the same in the two regions.

In future work, the wavelet-based model might be further refined by including multiple wavelets and accounting for eye movements. For instance, the current model assigns a single optimal wavelet (of all possible phases) to each neuron, potentially oversimplifying the diversity of receptive field structures. A better model might involve combinations of driving and suppressive Gabor wavelets (e.g., de Vries et al., 2020; Vintch et al., 2015). Another limitation is the absence of pupil tracking in our model, which could partly account for reduced repeatability, as saccades or undetected eye movements shift the visual input. A refinement of the model could account for the eye movements to help improve the reliability of feature estimation (Parker et al., 2023; Walker et al., 2019).

Furthermore, while the Zebra noise and analysis method that we used were optimized for mice, both stimulus and analysis can be readily adapted for species with higher spatial and temporal acuity, potentially including humans. We provide tools to generate custom libraries of Gabor wavelets, enabling finer feature descriptions, for instance in non-human primates. Likewise, our analytical framework is tailored for two-photon imaging but could be potentially adjusted to other modalities such as electrophysiological recordings, broadening its applicability to diverse experimental settings. Zebra noise could also be useful to map visual cortex with coarser methods, such as widefield imaging, functional ultrasound imaging, or fMRI. However, the analysis that we used assumes selectivity for orientation and spatial frequency, which is typically a poor assumption for the measurements obtained with coarse methods. The analysis would likely need to be modified by assuming the pooling of activity of multiple model neurons (Dumoulin & Wandell, 2008; Larsson & Heeger, 2006; Ribeiro et al., 2025).

In combination, the Zebra noise stimulus and the wavelet-based model provide a powerful means for rapid and robust ‘fingerprinting’ of neurons. This is highly valuable for tracking neuronal populations over time in studies of learning and plasticity. Furthermore, the model’s interpretability allows for direct comparison of tuning properties across visual regions, offering insights into the functional specialization of visual areas. This toolkit thus promises to accelerate progress in understanding how visual representations are organized, transformed by experience, and altered in disease.

## Acknowledgments

We thank Michael Krumin for help with the mesoscope, Charu Reddy for help with surgery, Ali Haydaroğlu for advice on data processing, Anyi Liu and Tinya Chang for testing our software, Cris Niell for the transgenic mouse line expressing soma-localized GCaMP8, and Patrick Reeson for helpful suggestions. This work was supported by EMBO (LTF 712-2021 to MS), the BRAIN Initiative (grant U01NS126057 to MC) and the Wellcome Trust (grant 223144/Z/21/Z to MC and KDH). MC holds the GlaxoSmithKline / Fight for Sight Chair in Visual Neuroscience.

## Author contributions

Contributions according to the *CRediT* taxonomy (https://credit.niso.org/).

|  | Carandini | Harris | Picard | Shinn | Skriabine |
| --- | --- | --- | --- | --- | --- |
| Conceptualization | • | • |  | • | • |
| Data curation |  |  | • |  | • |
| Formal analysis |  |  |  | • | • |
| Funding acquisition | • | • |  | • |  |
| Investigation |  |  | • |  | • |
| Methodology | • | • |  | • | • |
| Project administration | • |  |  |  |  |
| Software |  |  |  | • | • |
| Resources | • | • |  |  |  |
| Supervision | • | • |  |  |  |
| Validation |  |  |  |  | • |
| Visualization | • |  |  | • | • |
| Original draft | • |  |  | • | • |
| Review & editing | • | • |  | • | • |

## Methods

All procedures were conducted in accordance with the UK Animals Scientific Procedures Act (1986) under personal and project licenses released by the Home Office following appropriate ethics review.

### Zebra noise

Zebra noise is a comb-thresholded 3D Perlin noise. Below we describe the steps in generating it, and the analyses that we performed.

The first step is to generate 3D Perlin noise. Perlin noise is a type of gradient based noise widely used in computer graphics to generate texture or naturalistic terrains, and is attractive because it is fast to compute, and can be layered at different spatial and temporal scales to form a fractal 1/f-like spectrum. Notably, Perlin noise is computed on a pixel-by-pixel basis, allowing frames of the stimulus to be generated one-by-one with low memory requirements and processing power, permitting memory-efficient generation and parallel processing. By comparison, generating true 1/f pink noise (noise with a logarithmic frequency spectrum, from which Perlin Noise can be seen as a variation (Niell & Stryker, 2008)) quickly becomes computationally intractable due to the need to perform an inverse Fourier transform of a very large tensor.

To generate Perlin noise, an N-dimensional grid is generated. At each grid node, a random gradient vector is assigned. Then for each grid node we computed the dot product between the gradient vector and the node-point distance, for each point in the N-dimensional cells defined by the grid. The value is then interpolated across each value, resulting in a smooth random noise. This is performed at multiple spatial and temporal scales to produce a fractal spectrum. The process is performed in 3D so as to generate a movie.

The second step of Zebra noise generation is what we call a comb threshold, binning sub-intervals of the noise signal to generate sharp edge stripes. The Perlin noise values are rescaled to have a minimum value of 0 and a maximum value of 255 and then binned into k bins. We then assign the value of 1 to the odd bins and −1 to the even bins. This process creates a distinct high-contrast pattern of alternating sharp-edge stripes, from which Zebra noise derives its name. This texture exhibits both randomness and structured periodicity.

To measure the spatial autocorrelation of Zebra noise we computed the pairwise Pearson correlation of each pixel with a predetermined seed pixel at the center of the screen (**Figure 1e**). To analyze the spatial frequency and orientation content of Zebra noise (**Figure 1f**), we computed the power spectral density (PSD) of the stimulus images using the Fast Fourier Transform. We then transformed the PSD into polar coordinates to extract its radial (frequency) and angular (orientation) components. Finally, we computed the mean and standard deviation of the power at each frequency and orientation.

### Experiments

Experiments were conducted on 3 adult mice aged between ~3 weeks and 6 months. One mouse (female) expressed soma-localized GCaMp8m in excitatory neurons. It was obtained by crossing a tetO-RiboL1-jGCaMP8m transgenic mouse (generated by Denise Piscopo in the lab of Cris Niell at University of Oregon) with a Camk2a-tTA mouse (https://www.jax.org/strain/003010). The other two (one male and one female) expressed GCaMP6s in excitatory neurons. They were of C57BL/6 background and carried the wildtype allele of *Cdh23* that prevents age-related hearing loss (https://www.jax.org/strain/002756). They were obtained by crossing a TetO-GCaMP6s-Cdh23 line (https://www.jax.org/strain/024742) (Wekselblatt et al., 2016) with a Camk2a-tTA-Cdh23 mouse (https://www.jax.org/strain/003010).

### Surgical procedure

For the implant surgery, mice were anesthetized with isoflurane (5% for induction and 0.5–1% during the procedure) and secured in a stereotaxic frame. An analgesic, Rimadyl (5 mg/kg), was administered subcutaneously prior to the procedure and orally for the three following days. To prevent brain edema Dexamethasone (0.5 mg/kg, IM) was administered intramuscularly 30 minutes prior to surgery. Throughout the procedure, the exposed brain was continuously perfused with artificial cerebrospinal fluid (150 mM NaCl, 2.5 mM KCl, 10 mM HEPES, 2 mM CaCl2, 1 mM MgCl2; pH 7.3 adjusted with NaOH, 300 mOsm). A headplate was implanted for subsequent head-fixation, and a craniotomy (3 to 4 mm in diameter) was performed to implant a cranial window for optical access, made of three-layer sandwiched borosilicate glass. After they recovered (minimum of 3 days after the surgery, in practice, 1 week), mice were habituated to the rig and trained to be head fixed.

### Two-photon imaging

We performed two-photon imaging using a standard resonant mesoscope (Sofroniew et al., 2016) (Multiphoton Mesoscope, ThorLabs Inc) controlled by ScanImage 4.2 (Pologruto et al., 2003). Excitation light was delivered by a 920 nm ultrafast laser (FemtoFibre ultra 920, Toptica Photonics AG, Graefelfing, Germany). Imaging depth was adjusted using a remote focusing mirror with 1 mm travel range. Fluorescent signal was collected in the green channel (Semrock FF01-520/70 emission filter) by a GaAsP PMT (PMT2103, Thorlabs, H10770PA-40, Hamamatsu Photonics), amplified by a fast transimpedance preamplifier (400 MHz band-width, HCA-400M-5K-C, FEMTO Messtechnik GmbH, Berlin, Germany), and digitized by a vDAQ acquisition board (Vidrio Technologies, MBF Bioscience, Williston, VT, USA). Acquisition sampling was synchronized to the laser pulses.

To prevent visual stimulation light from contaminating the fluorescence signal, the monitor backlight was rapidly switched on/off by a custom circuit implemented on Teensy 4.0, such that the monitor light was on only when the resonant mirror was making a U-turn and no image-building fluorescence was collected. In addition, to prevent the PMT from saturating or tripping, the imaging objective was light-shielded using a custom-made plastic cone.

Fields of view (FOV), roughly centered on V1, were imaged with a resolution of 2097× 1536 pixels at 3-4 Hz. FOVs typically spanned 3 stripes of size 2097 × 512 pixels covering a window of 2867 × 2100μm for the imaging of neurons in a single plane in V1 L2/3. The acquisition resolution was 1.3671 μm per pixel.

The raw neural images were processed with Suite2p (Pachitariu et al., 2017) to extract the firing rate of the neuron as a time series. The firing rates were upsampled to 30 Hz to match the stimulus frame rate.

### Traditional visual stimuli

Visual stimuli were displayed at 30 Hz on two screens placed at 90 degrees in front and on the left side of the head-fixed animal, covering the range of −135 to 45 deg of azimuth and −35 to 35 deg of elevation. Experiments were 3 hours long, and each session included 10 min of spontaneous activity, 3 × 10 min of sparse noise, 3 × 18 directions drifting gratings, and 3 × 7 min of Zebra noise.

To map the early retinotopic organization of the mouse visual cortex, we used a sparse noise stimulus consisting of black and white squares presented on a gray background. Individual squares, each measuring 9 × 9 degrees of visual angle, appeared randomly at different positions on the two screens. The stimulus was presented for a total duration of 10 minutes, ensuring sufficient sampling of receptive fields across the visual space.

To assess neuronal tuning to orientation and direction, we presented full-field drifting gratings spanning both monitors. The gratings were shown in 18 evenly spaced orientations (10° increments) with a randomly assigned forward or backward drift. Each grating was presented five times, with a duration of 1 s, followed by a 1-s inter-stimulus interval with a gray screen. The gratings had a spatial frequency of 0.05 cycles per degree and a drift rate of 2 Hz.

We also performed a separate experiment to measure responses to the Trippy stimulus (MICrONS Consortium et al., 2025). The associated code (Yatsenko et al., 2018) produced up to a few minutes of stimulus. We thus ran the stimulus for 1 minute (3 repeats). For fairness, we compared those responses to Zebra noise that was run for the same duration.

### Wavelet-based model

To characterize neuronal selectivity with high spatiotemporal precision, we developed a wavelet-based approach that decomposes any visual stimuli into a structured library of localized wavelets. The code for this analysis is contained in a package called WavEn, which contains a graphical user interface (Supplementary Figure 9). The package is available on GitHub, at https://github.com/skriabineSop/waven.

### Gabor library and filtering

To extract spatial features from the visual stimulus, we constructed two libraries of Gabor wavelets. Each wavelet is defined by a center location (azimuth between −45 and 135 deg, elevation between –35 and 35 deg), an orientation (8 evenly spaced angles between 0° and 180°), a spatial frequency, a size (defined as Gaussian envelope radius at half maximum), and a phase (sine or cosine). We assembled these wavelets in two libraries, one coarse and one fine. The *coarse* library contains 28,512 wavelets (27 azimuths × 11 elevations × 8 orientations × 6 size/frequency combinations × 2 phases). In this library, spatial frequency and size are linearly coupled, enabling rapid estimation of neuronal receptive field position and orientation with a compact set of wavelets. The *fine* library contains 2,332,800 wavelets (135 azimuths × 54 elevations × 8 orientations × 5 sizes × 4 frequencies × 2 phases), with spatial frequency and size sampled independently. This library allows more refined modeling of visual properties, including precise receptive field location and tuning for scale and frequency.

The raw stimulus displayed on the screen during the experiment had high resolution (1200 × 600 pixels). For processing time, it was downsampled to 135 × 54 pixels before applying the wavelet transform, which allows to reduce the number of Gabor wavelets. Each frame of the movie was then filtered using the dot product with every wavelet in the library.

### Step 1: Coarse feature estimation

We first obtained a rough estimate of a neuron’s receptive field properties using the coarse Gabor wavelets. For each wavelet, we filtered the Zebra noise video with the wavelet, as in **Figure 3**. Because each wavelet is defined at two phases (0° and 90°), we applied the filtering separately for each phase and then summed the squared outputs. This yielded for each wavelet, one vector that can be interpreted as the variation of the wavelet amplitude. We then computed the Pearson’s correlation these timeseries and each neuron’s deconvolved and z-scored activity. This procedure provided one correlation value per wavelet, serving as a measure of similarity between the neuron, and the expected activity of the neuron and a model complex neuron described by that particular wavelet.

### Step 2: Refined feature estimation

Step 2 aims to refine the parameters (x, y, θ, s, f) of a neuron (**Figure 4a**). We refined our initial coarse estimates of neuron features using our fine feature Gabor wavelets. Due to the large number of parameters of the fine Gabor wavelets, computational runtime prevented us from running the same procedure as with the coarse Gabor wavelets. The fine Gabor wavelets provide essential information about neurons, in-cluding the phase, the ratio between spatial frequency and receptive field size, and a high-resolution estimate of receptive field location. Therefore, we designed an algorithm to efficiently estimate these properties using the fine Gabor wavelet library. By iteratively fixing one parameter at a time (e.g., position) and refining the others (e.g., orientation, frequency, scale), we progressively improve each neuron’s receptive field position estimate as much as possible, within the limits of the filter library. The procedure is detailed below:

1. Initialize position (x0, y0) based on the earlier coarse estimation
2. Compute Pearson correlation of all fine wavelets within (x0±dx, y0±dy) with neuron activity for both phases separately. Select the wavelet with the highest correlation score (across both phases) in absolute value. Fix a new optimal position x, y accordingly.
3. Estimate initial orientation (o), frequency (f), and scale (s)
4. While not converged
  - Fix orientation (o), frequency (f), and scale (s)
  - Choose the wavelet with position (x, y) which maximizes Pearson correlation
  - Fix position (x, y)
  - Choose the orientation (o), frequency (f), and scale (s) which maximize Pearson correlation

Using the resulting correlations, we compute the tuning curve for each of these parameters. We define the tuning curve for each model parameter x, y, θ, s, f to be the correlation of the neuron’s activity with the wavelets where that specific parameter is varied, and all other parameters are optimal (**Figure 4f**). This is analogous to predicting the response to experimental stimuli which probe a single parameter at a time, such as varying the direction of drifting gratings.

Step 3: Model fitting

Step 3 aims to refine the estimate of each neuron’s receptive field by considering its preferences for phase (*φ*), drift (*φ*′) and amplitude (*A*)

Let us call {*w*_1_, *w*_2_, …, *w*_*W*_} the wavelets in a library. Let’s assume our neuron’s best-matching wavelet (as found in Step 2) is *w*_*j*_. This wavelet *w*_*j*_ is a matrix of complex numbers whose real part corresponds to the 0° phase an imaginary part corresponds to the 90° phase. Let *p*_*j*_(*t*) be the projection of stimulus frame *t*on the wavelet *w*_*j*_. It is a complex vector whose real and imaginary components indicate projections on the wavelets with 0° and 90° phase. We then compute the magnitude *A*(*t*) = |*p*_*j*_(*t*)|and phase *φ*(*t*) = *arg* (*p*_*j*_(*t*)) of the wavelet-transformed stimulus, as well as the derivative of the phase *φ*(*t*)′ = *dφ* (*t*)/*dt*. Based on these three prop-erties, *α*(*t*), *φ*(*t*), and *φ*′(*t*), we build a 3D weighted histogram of the firing rate of the neuron, and its 3D occupancy matrix for these three properties (**Figure 4d** and Supplementary Figure 4). (20 bins per dimension). To improve robustness and to extrapolate to regions not covered by the stimulus, we blurred the 3D matrix of firing rates of the neuron and the 3D occupancy matrix separately before dividing the two. A similar procedure is typically used to compute two-dimensional place fields (O’ Keefe & Burgess, 1996). To evaluate the model, we used the resulting 3D matrix as a lookup table: for any combination of phase, amplitude and drift: *φ*(*t*),*A*(*t*), and *φ*′(*t*), we estimated the predicted firing rate using nearest-neighbor interpolation into the 3D matrix. This step allows us to approximate the expected response for arbitrary tuples, even if they do not fall exactly on bin centers. To extract phase, drift, and amplitude tuning curves (**Figure 4g**), we average the response matrix along two of its three dimensions, each time isolating the dimension of interest.

### Cross-validation and performance metrics

The model predictions are cross-validated as follows: the model estimates the different tuning from the repeats number 1 and 3, and the correlation of the prediction is computed from the remaining repetitions (Figure 4 and 5).

To assess the model’s ability to generalize to previously unseen stimuli, we reserved the last 1 min of the stimulus as a test set, while training was performed only on the first 1-6 min of stimulus.

We define repeatability as the Pearson correlation between the mean responses computed from two subsets of stimulus repetitions of equal size. This measure is equivalent to the performance of an “oracle” (Vintch et al., 2015) and to the fraction of explainable variance (Du et al., 2025). We define predictability as the Pearson correlation between the predicted firing rate and the neuron’s actual firing rate, computed as the average activity on the test set. We define the prediction score as the ratio of predictability to repeatability.

## Additional analyses

### Smooth estimates of preferred position

When generating sign maps, it is necessary to compute retinotopic gradients (Sereno et al., 1995). To do so, we must impose spatial smoothness on the retinotopic representation and disregard the local scatter observed at the single-neuron scale. For this purpose, when computing sign maps, we smoothed the estimated receptive field positions of the neurons by taking the median of the estimated position for neighboring neurons within a 25 μm radius. This step is optional and can be useful to generate retinotopic maps and segment visual areas. To avoid bias, however, we advise not to use it when analyzing the properties of individual neurons.

### Sign maps and visual areas

Using each neuron’s preferred retinotopic position and anatomical coordinates, we interpolated these values to estimate the preferred azimuth and elevation at every pixel across the im-aging plane, thereby generating continuous retinotopic maps. These maps were then smoothed using a Gaussian kernel to ensure gradient continuity and suppress local noise. We then extract the visual field sign at each pixel, computed as the sine of the angle between the local gradients in azimuth and elevation (Sereno et al., 1995, Zhuang et al., 2017).

We use the sign map’s regions to segment the cortical space into distinct zones, allowing us to categorize neurons based on the location of their receptive fields in the visual field.: From the sign map, we performed image segmentation using a modified watershed algorithm to identify distinct regions in the visual cortex from their signed values. First, a binary mask was generated based on the signed map, where the threshold was determined by the sign of the values. A morphological opening operation was applied to remove small noise in the binary map. Each pixel was assigned to be “sure background”, “sure foreground”, or “unknown”. To define the regions, the sure background was obtained by dilating the binary map, while the sure foreground was determined using the distance transform followed by a thresholding. The unknown region was then calculated by subtracting the sure foreground from the sure background. Connected components in the sure foreground were labeled to form markers for the watershed algorithm. The markers were adjusted by incrementing the labels of the sure background and setting the unknown regions to zero. The watershed algorithm was then applied to segment the regions, with the signed map serving as the input for the merging process.

Finally, for each neuron we identified its corresponding visual area by extracting the watershed labels at its position on the imaging plane, allowing us to segment and classify neuronal regions in the image. V1 was nicely segmented. We define as ‘visually responsive neurons’ all neurons with a repeatability higher than 0.2. We define as Higher visual areas (HVA) any neurons that do not belong to V1 but are visually responsive.

## Supplementary Figures

**Supplementary Figure 1.**
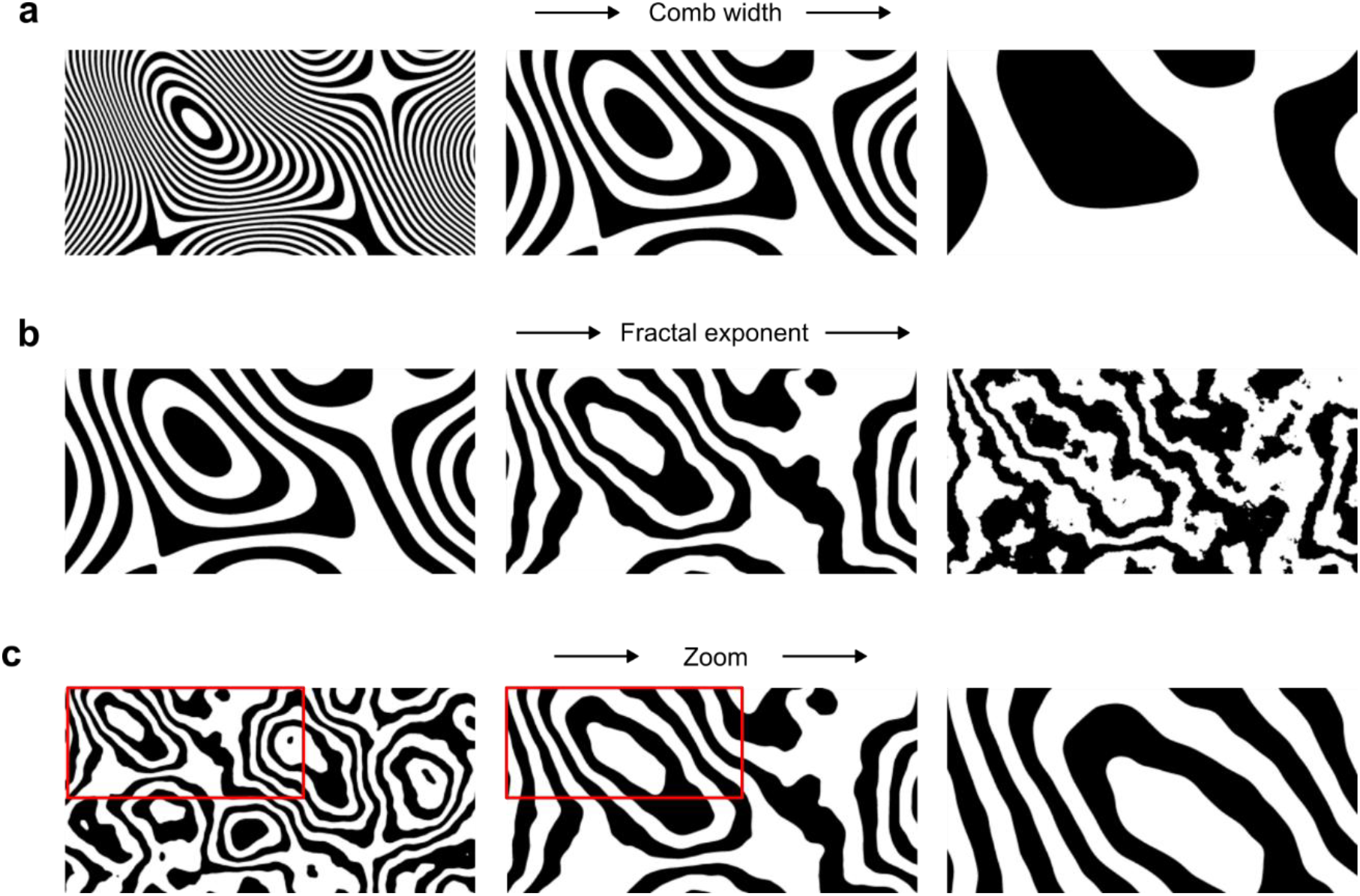
Parametrization of Zebra noise. The Zebra noise stimulus is controlled by three parameters: comb width, fractal exponent and zoom magnitude. **a:** Effect of increasing comb width. **b**, Effect of increasing fractal exponent. **c**, Effect of increasing zoom magnitude.

**Supplementary Figure 2.**
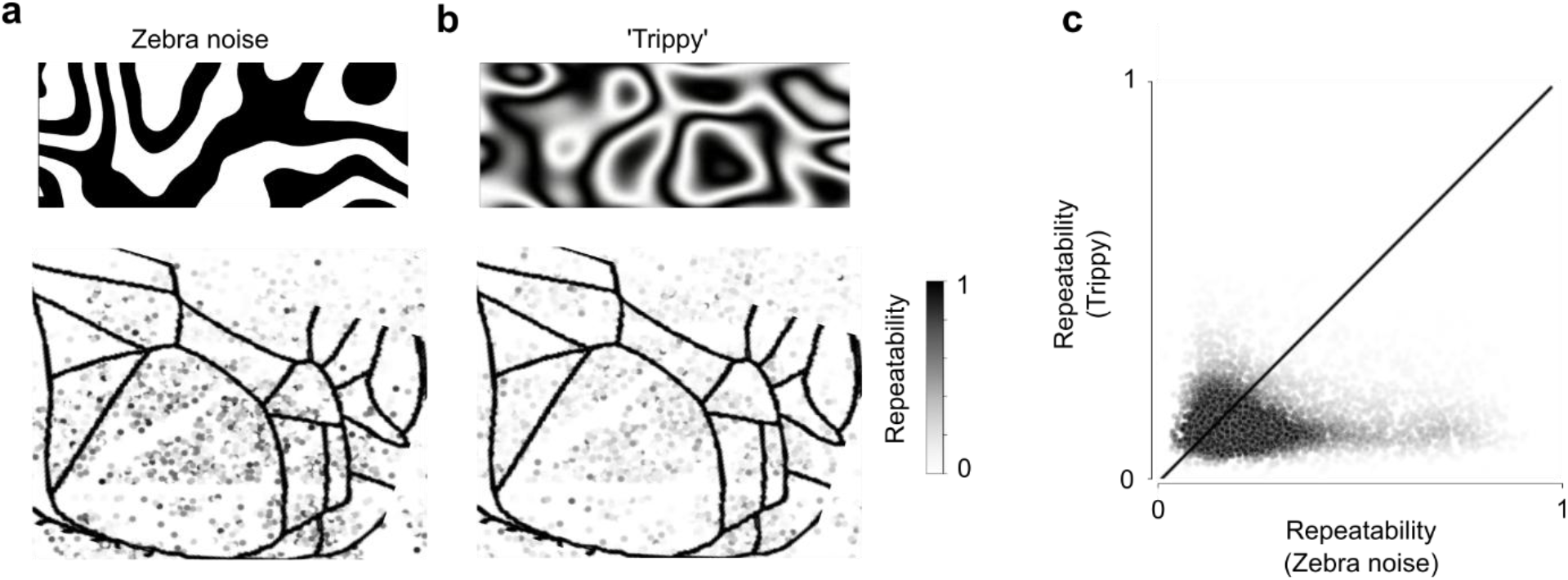
Zebra noise elicits more repeatable neuron responses than Trippy. Short snippets of Zebra noise and Trippy noise (3 repeats of a 1 min stimulus) were presented to a mouse expressing GCaMP6s in excitatory neurons. **a**: Repeatability (average Pearson’s correlation of neural responses between trials) of the responses to Zebra noise, expressed as gray level. Each dot corresponds to a neuron. **b:** Repeatability of the responses to Trippy noise. **c:** Comparison of repeatability to Zebra noise (abscissa) vs. Trippy noise (ordinate). Each dot is a neuron. In most neurons, Zebra noise elicited more repeatable responses than Trippy noise. This increased repeatability may be due to the presence of sharp edges in Zebra noise.

**Supplementary Figure 3.**
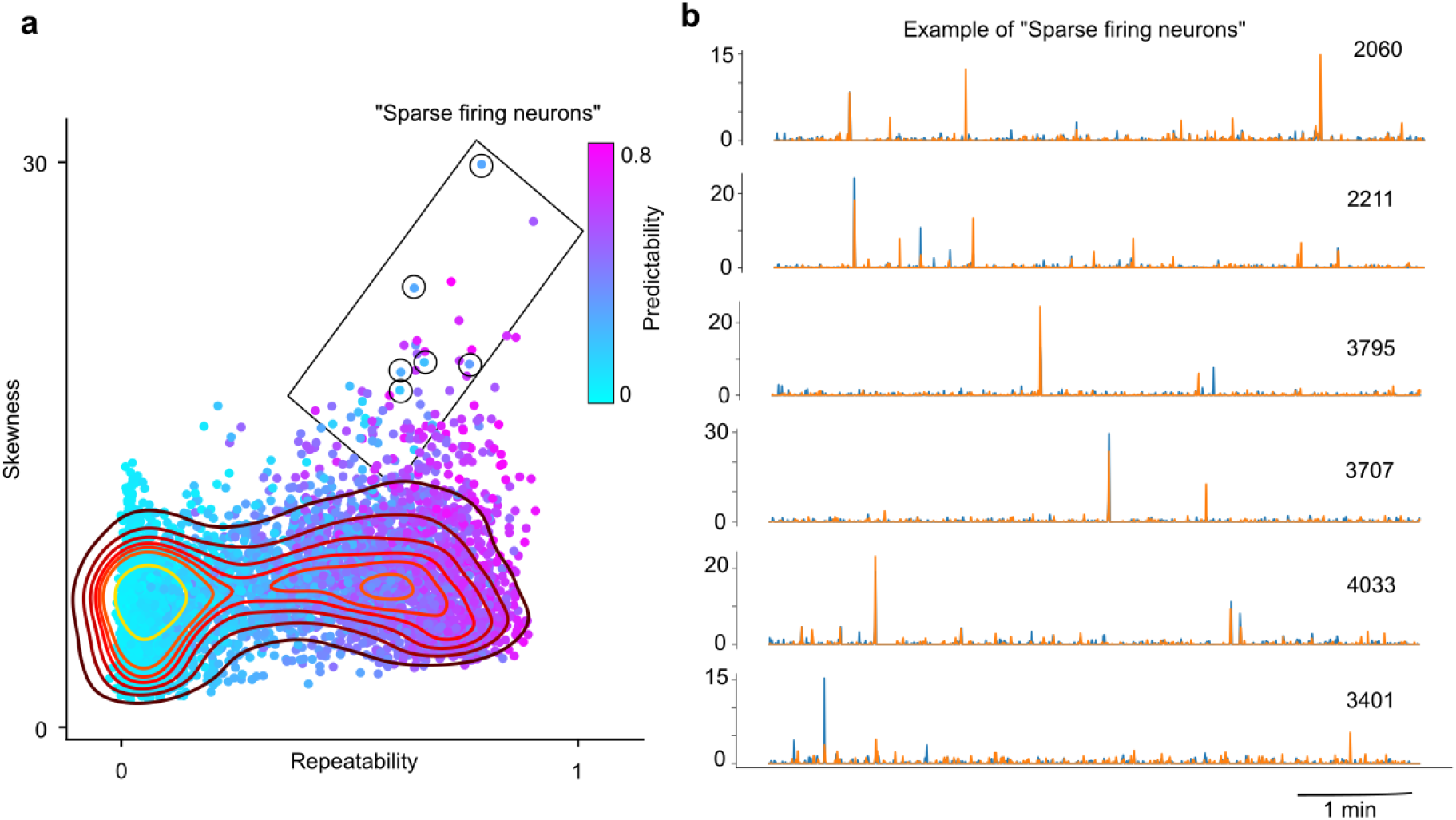
The wavelet-based model can capture the responses of neurons that fire sparsely. **a**: Skewness vs. repeatability for 4,558 neurons recorded from the visual cortex and the surrounding regions (same experiment as **Figure 4**). The color indicates the predictability by the wavelet-based model. Circles highlight 6 cells with high repeatability, high skewness, and low predictability. **b**, Responses of those 6 example cells on repeat 1 and 3 (*blue*) and repeat 2 and 4 (*orange*).

**Supplementary Figure 4.**
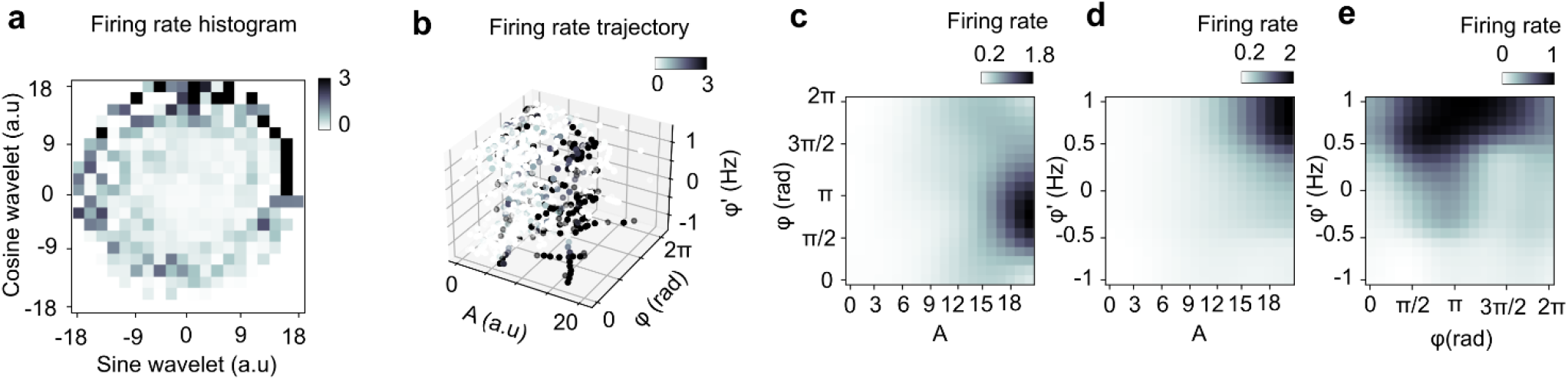
Mapping the selectivity for amplitude, phase, and drift rate. a: Firing rate of the neuron in Figure 4 expressed as a function of the sine and the cosine projection of its corresponding best fitting wavelet. b: Firing rate as a function of amplitude *A*, phase *φ*, and drift rate *φ*′. c: Firing rate as a function of phase and amplitude. d: Firing rate as a function of amplitude and drift rate. e: Firing rate as a function of phase and drift rate. Grayscale represents the firing rate (insets).

**Supplementary Figure 5.**
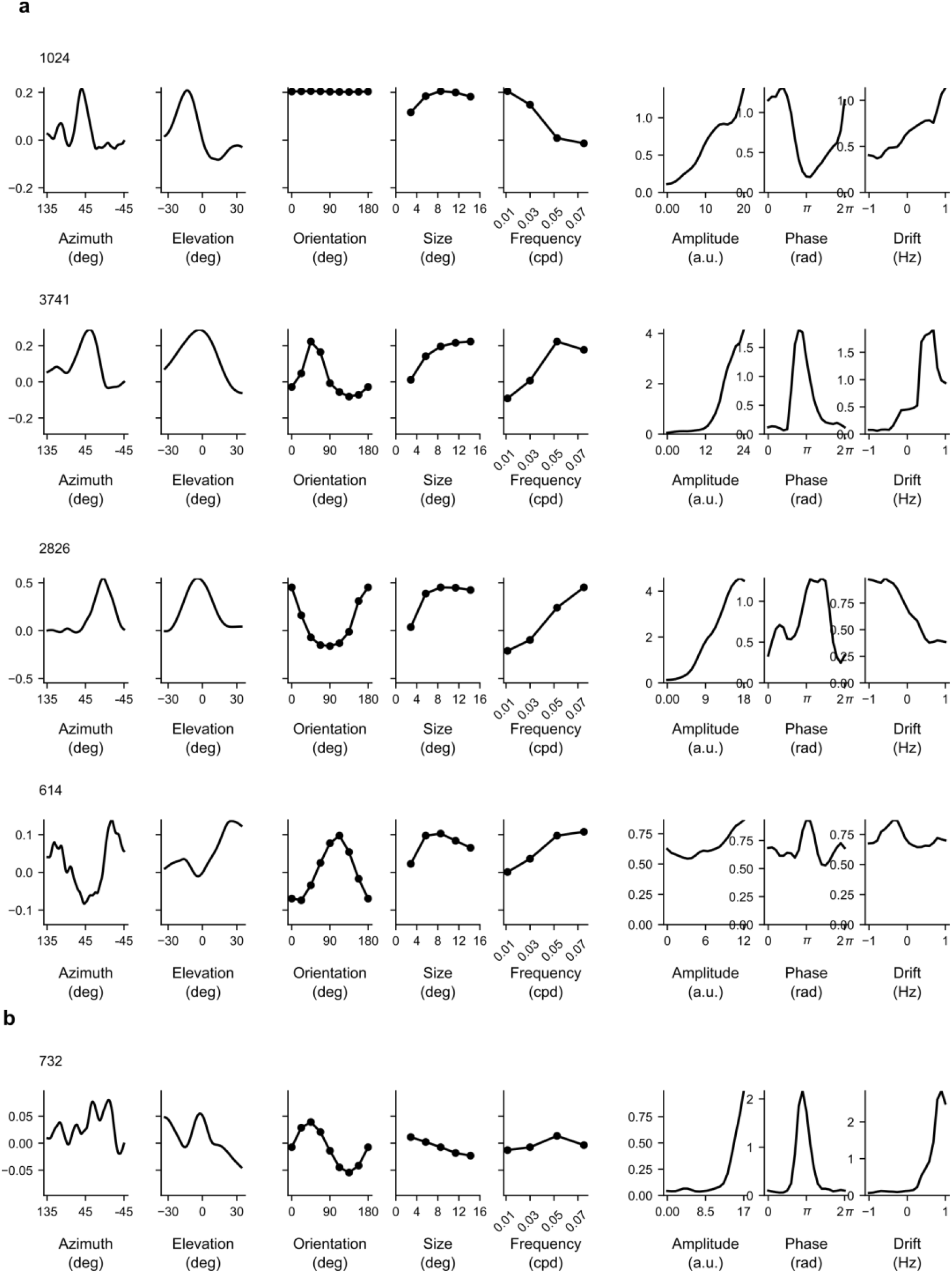
Examples of neuron tuning curves measured with Zebra noise and produced with the wavelet-based model. Each row shows the results for a given neuron, using the same format as in Figure 4f,g. a: Examples of cells tuned for different combinations of orientation, size, and spatial frequency. b: Example of a cell with ambiguous receptive field location.

**Supplementary Figure 6.**
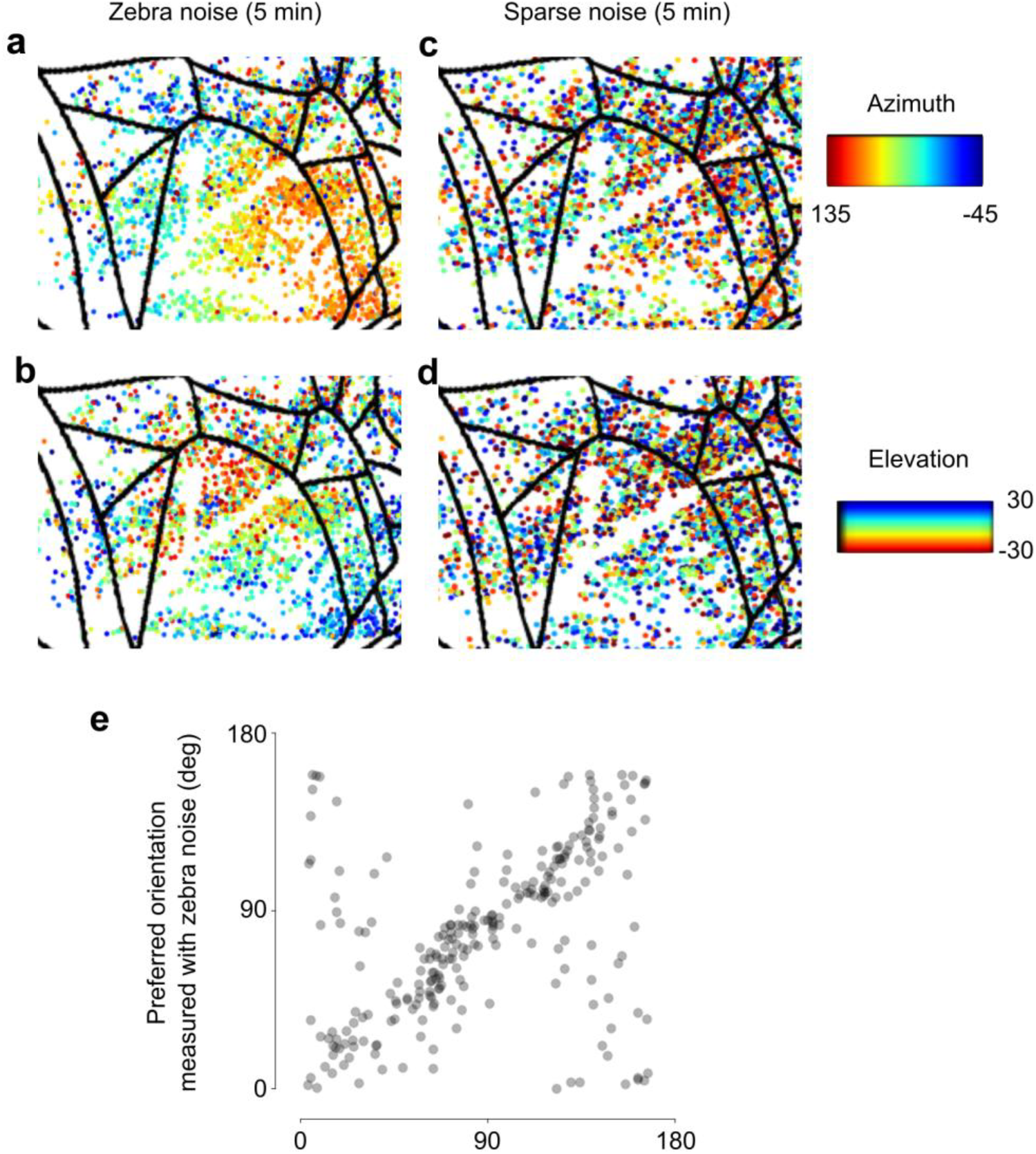
Zebra noise yields better retinotopic maps than sparse noise and similar orientation tuning as gratings. **a,b:** Retinotopy measured with Zebra noise, showing selectivity for azimuth (a) and elevation (b). Each dot represents a neuron. **c,d**: Same, measured with sparse noise (black and white squares randomly flashed on the screen). In both cases, stimuli were 5 min long and were repeated 3 times. **e:** Comparison of orientation preference computed with Zebra noise (*ordinate*) and drifting gratings (*abscissa*). Each dot is a neuron. Only neurons with circular variance <0.3 were included in this analysis.

**Supplementary Figure 7.**
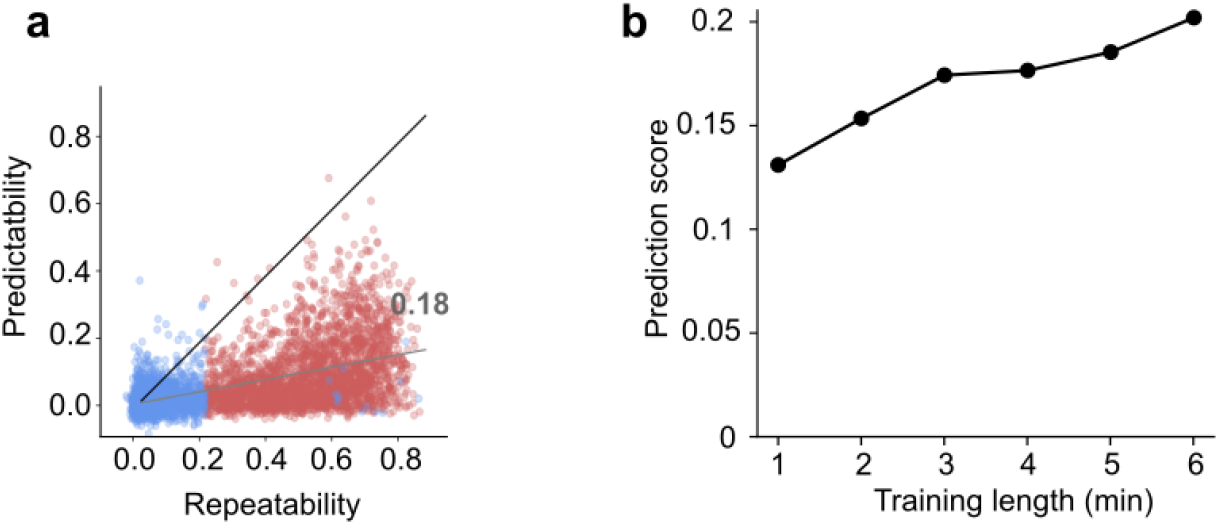
Ability of the model to predict responses in an unseen stimulus (generalizability). Data from the same dataset as in Figure 5. **a**: Performance of the model when the training data is obtained from 5 minutes of stimulus and the test data are obtained from a different 1 minute of stimulus, as a function of repeatability in the test period. **b:** Prediction score (ratio of predictability to repeatability) as a function of the length of the training set, when training and testing are done on a different 1-min stimulus sequence.

**Supplementary Figure 8.**
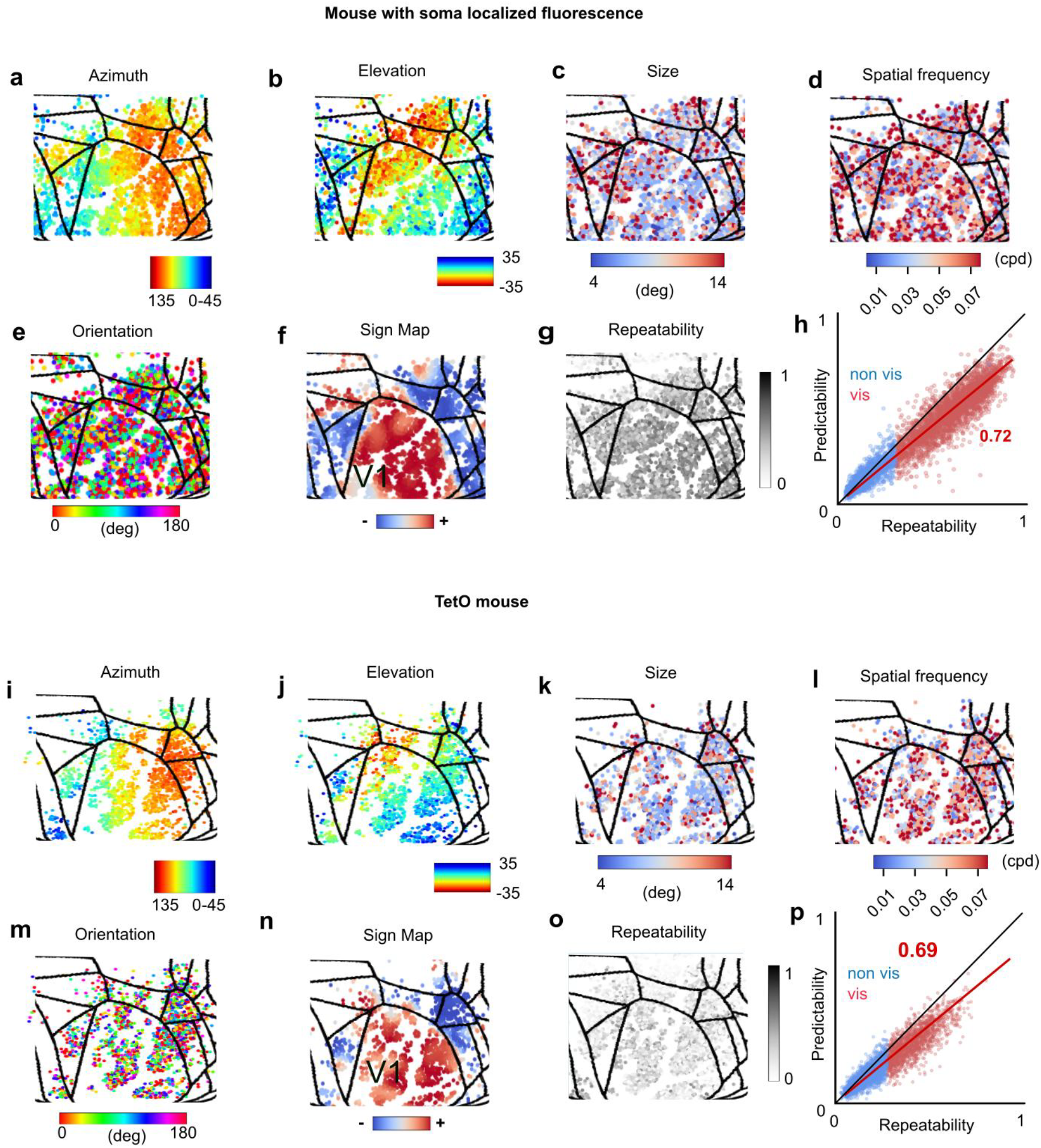
Data obtained in two additional mice. For each mouse, we show data in the same format as **Figure 5. a-h:** Results in a mouse expressing soma-localized GCaMP8m. **i-p:** Results for a mouse expressing GCaMP6s (same line as in **Figure 5**).

**Supplementary Figure 9.**
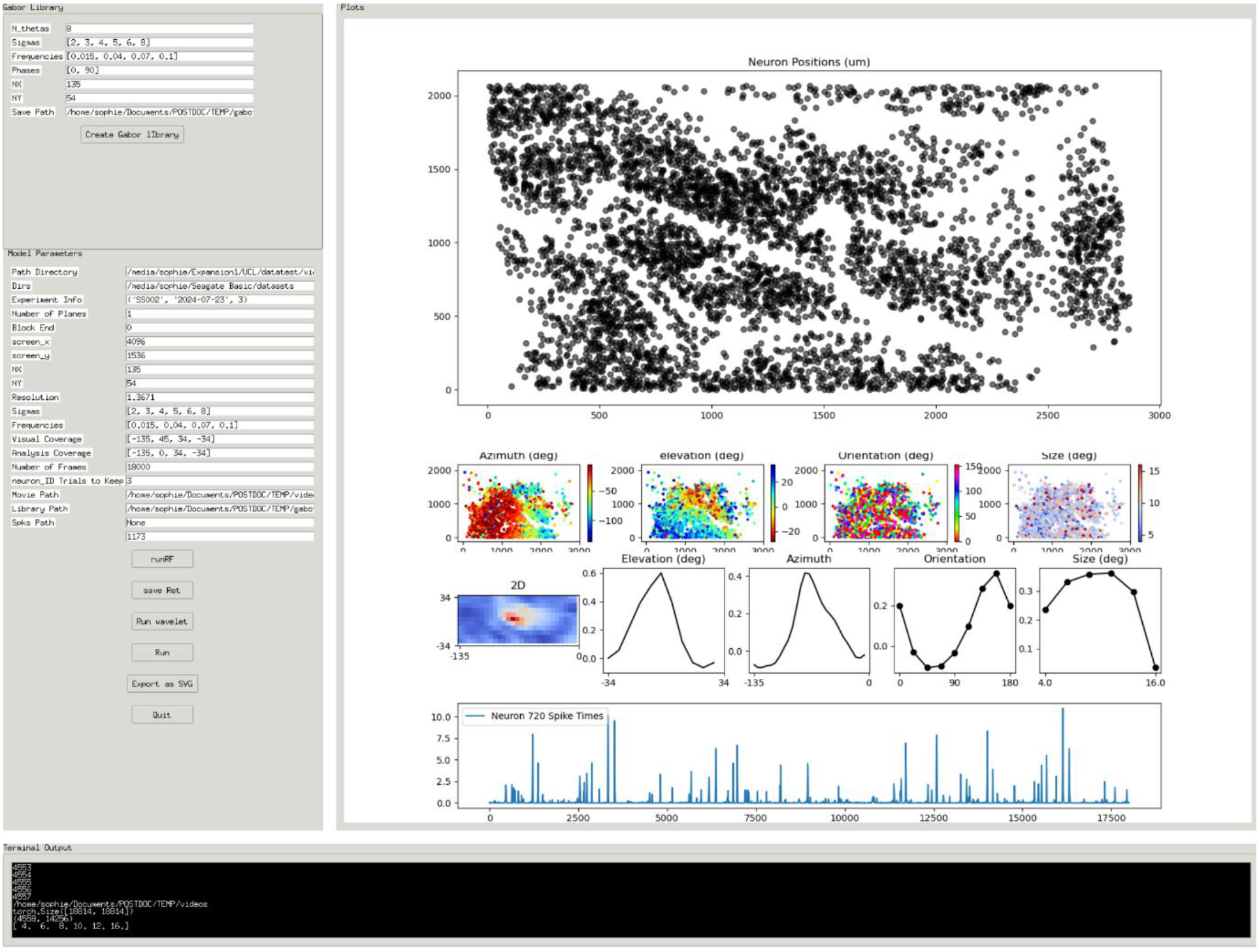
Overview of the Graphical User Interface. The left panel displays all configurable parameters. Default values are provided and are optimized for single-plane mesoscope experiments, though they should be suitable for most use cases. Further details about these parameters are available on the GitHub page (https://github.com/skri-abineSop/waven). The right panel shows the output plots, from top to bottom: a scatter plot of neuron positions; maps for azimuth, elevation, orientation, and scale (restricted to visually responsive neurons); map of the receptive field location; tuning curves; and the spike time series.

## References

Benucci, A., Ringach, D. L., & Carandini, M. (2009). Coding of stimulus sequences by population responses in visual cortex. Nature Neuroscience, 12(10), 1317–1324. 10.1038/nn.2398

Bonin, V., Histed, M. H., Yurgenson, S., & Reid, R. C. (2011). Local Diversity and Fine-Scale Organization of Receptive Fields in Mouse Visual Cortex. Journal of Neuroscience, 31(50), 18506–18521. 10.1523/JNEUROSCI.2974-11.2011

Cadena, S. A., Denfield, G. H., Walker, E. Y., Gatys, L. A., Tolias, A. S., Bethge, M., & Ecker, A. S. (2019). Deep convolutional models improve predictions of macaque V1 responses to natural images. PLOS Computational Biology, 15(4), e1006897. 10.1371/JOURNAL.PCBI.1006897

de Vries, S. E. J., Lecoq, J. A., Buice, M. A., Groblewski, P. A., Ocker, G. K., Oliver, M., Feng, D., Cain, N., Ledochowitsch, P., Millman, D., Roll, K., Garrett, M., Keenan, T., Kuan, L., Mihalas, S., Olsen, S., Thompson, C., Wakeman, W., Waters, J., … Koch, C. (2020). A large-scale standardized physiological survey reveals functional organization of the mouse visual cortex. Nature Neuroscience, 23(1), 138– 151. 10.1038/s41593-019-0550-9

Dhruv, N. T., & Carandini, M. (2014). Cascaded effects of spatial adaptation in the early visual system. Neuron, 81(3), 529–535. 10.1016/j.neuron.2013.11.025

Dräger, U. C. (1975). Receptive fields of single cells and topography in mouse visual cortex. The Journal of Comparative Neurology, 160(3), 269–289. 10.1002/CNE.901600302

Du, F., Angel Núñez-Ochoa, M., Pachitariu, M., & Stringer, C. (2025). A simplified minimodel of visual cortical neurons. Nature Communications 2025 16:1, 16(1), 1–13. 10.1038/s41467025-61171-9

Dumoulin, S. O., & Wandell, B. A. (2008). Population receptive field estimates in human visual cortex. NeuroImage, 39(2), 647–660. 10.1016/J.NEUROIMAGE.2007.09.034

Erisken, S., Vaiceliunaite, A., Jurjut, O., Fiorini, M., Katzner, S., & Busse, L. (2014). Effects of locomotion extend throughout the mouse early visual system. Current Biology : CB, 24(24), 2899–2907. 10.1016/J.CUB.2014.10.045

Felsen, G., Touryan, J., Han, F., & Dan, Y. (2005). Cortical Sensitivity to Visual Features in Natural Scenes. PLOS Biology, 3(10), e342. 10.1371/JOURNAL.PBIO.0030342

Hubel, D. H., & Wiesel, T. N. (1959). Receptive fields of single neurones in the cat’s striate cortex. The Journal of Physiology, 148(3), 574–591. 10.1113/jphysiol.1959.sp006308

Jones, J. P., & Palmer, L. A. (1987a). An evaluation of the two-dimensional Gabor filter model of simple receptive fields in cat striate cortex. Journal of Neurophysiology, 58(6), 1233–1258. 10.1152/JN.1987.58.6.1233

Jones, J. P., & Palmer, L. A. (1987b). The two-dimensional spatial structure of simple receptive fields in cat striate cortex. Https://Doi.Org/10.1152/Jn.1987.58.6.1187, 58(6), 1187–1211. 10.1152/JN.1987.58.6.1187

Larsson, J., & Heeger, D. J. (2006). Two Retinotopic Visual Areas in Human Lateral Occipital Cortex. Journal of Neuroscience, 26(51), 13128–13142. 10.1523/JNEUROSCI.165706.2006

MICrONS Consortium, Bae, J. A., Baptiste, M., Bishop, C. A., Bodor, A. L., Brittain, D., Buchanan, J., Bumbarger, D. J., Castro, M. A., Celii, B., Cobos, E., Collman, F., Maçarico Da Costa, N., Dorkenwald, S., Elabbady, L., Fahey, P. G., Fliss, T., Froudarakis, E., Gager, J., … Zhang, C. (2023). Functional connectomics spanning multiple areas of mouse visual cortex. 10.1101/2021.07.28.454025

Movshon, J. A., Thompson, I. D., & Tolhurst, D. J. (1978a). Receptive field organization of complex cells in the cat’s striate cortex. The Journal of Physiology, 283(1), 79–99. 10.1113/JPHYSIOL.1978.SP012489

Movshon, J. A., Thompson, I. D., & Tolhurst, D. J. (1978b). Spatial summation in the receptive fields of simple cells in the cat’s striate cortex. The Journal of Physiology, 283(1), 53–77. 10.1113/jphysiol.1978.sp012488

Niell, C. M., & Stryker, M. P. (2008). Highly Selective Receptive Fields in Mouse Visual Cortex. Journal of Neuroscience, 28(30), 7520–7536. 10.1523/JNEUROSCI.062308.2008

Niell, C. M., & Stryker, M. P. (2010). Modulation of visual responses by behavioral state in mouse visual cortex. Neuron, 65(4), 472–479. 10.1016/j.neuron.2010.01.033

Ohki, K., & Reid, R. C. (2007). Specificity and randomness in the visual cortex. Current Opinion in Neurobiology, 17(4), 401–407. 10.1016/J.CONB.2007.07.007

O’ Keefe, J., & Burgess, N. (1996). Geometric determinants of the place fields of hippocampal neurons. Nature 1996 381:6581, 381(6581), 425–428. 10.1038/381425a0

Pachitariu, M., Stringer, C., Dipoppa, M., Schröder, S., Rossi, L. F., Dalgleish, H., Carandini, M., & Harris, K. D. (2017). Suite2p: beyond 10,000 neurons with standard two-photon microscopy. BioRxiv, 061507. 10.1101/061507

Parker, P. R. L., Martins, D. M., Leonard, E. S. P., Casey, N. M., Sharp, S. L., Abe, E. T. T., Smear, M. C., Yates, J. L., Mitchell, J. F., & Niell, C. M. (2023). A dynamic sequence of visual processing initiated by gaze shifts. Nature Neuroscience, 26(12), 2192– 2202. 10.1038/S41593-023-01481-7

Perlin, K. (1985). An image synthesizer. SIGGRAPH Comput. Graph., 19(3), 287–296. 10.1145/325165.325247

Pologruto, T. A., Sabatini, B. L., & Svoboda, K. (2003). ScanImage: Flexible software for operating laser scanning microscopes. BioMedical Engineering OnLine, 2, 13. 10.1186/1475-925X-2-13

Ribeiro, F. L., Benson, N. C., & Puckett, A. M. (2025). Human retinotopic mapping: From empirical to computational models of retinotopy. Journal of Vision, 25(8), 14. 10.1167/JOV.25.8.14

Ringach, D. L., Hawken, M. J., & Shapley, R. (1997). Dynamics of orientation tuning in macaque primary visual cortex. Nature, 387(6630), 281–284. 10.1038/387281a0

Schröder, S., Steinmetz, N. A., Krumin, M., Pachitariu, M., Rizzi, M., Lagnado, L., Harris, K. D., & Carandini, M. (2020). Arousal Modulates Retinal Output. Neuron, 107(3), 487-495.e9. 10.1016/J.NEURON.2020.04.026

Sereno, M. I., Dale, A. M., Reppas, J. B., Kwong, K. K., Belliveau, J. W., Brady, T. J., Rosen, B. R., & Tootell, R. B. H. (1995). Borders of multiple visual areas in humans revealed by functional magnetic resonance imaging. Science (New York, N.Y.), 268(5212), 889–893. 10.1126/SCIENCE.7754376

Sofroniew, N. J., Flickinger, D., King, J., & Svoboda, K. (2016). A large field of view two-photon mesoscope with subcellular resolution for in vivo imaging. ELife, 5(JUN2016). 10.7554/ELIFE.14472

Tong, R., Silva, R. da, Lin, D., Ghosh, A., Wilsenach, J., Cianfarano, E., Bashivan, P., Richards, B., & Trenholm, S. (2023). The feature landscape of visual cortex. BioRxiv, 2023.11.03.565500. 10.1101/2023.11.03.565500

Touryan, J., Felsen, G., & Dan, Y. (2005). Spatial structure of complex cell receptive fields measured with natural images. Neuron, 45(5), 781–791. 10.1016/J.NEURON.2005.01.029

Vintch, B., Movshon, J. A., & Simoncelli, E. P. (2015). A Convolutional Subunit Model for Neuronal Responses in Macaque V1. Journal of Neuroscience, 35(44), 14829–14841. 10.1523/JNEUROSCI.281513.2015

Walker, E. Y., Sinz, F. H., Cobos, E., Muhammad, T., Froudarakis, E., Fahey, P. G., Ecker, A. S., Reimer, J., Pitkow, X., & Tolias, A. S. (2019). Inception loops discover what excites neurons most using deep predictive models. Nature Neuroscience, 22(12), 2060–2065. 10.1038/s41593-0190517-x

Wang, E. Y., Fahey, P. G., Ding, Z., Papadopoulos, S., Ponder, K., Weis, M. A., Chang, A., Muhammad, T., Patel, S., Ding, Z., Tran, D., Fu, J., Schneider-Mizell, C. M., da Costa, N. M., Reid, R. C., Collman, F., da Costa, N. M., Franke, K., Ecker, A. S., … Tolias, A. S. (2025). Foundation model of neural activity predicts response to new stimulus types. Nature 2025 640:8058, 640(8058), 470–477. 10.1038/s41586-025-08829-y

Wang, Q., Ding, S. L., Li, Y., Royall, J., Feng, D., Lesnar, P., Graddis, N., Naeemi, M., Facer, B., Ho, A., Dolbeare, T., Blanchard, B., Dee, N., Wakeman, W., Hirokawa, K. E., Szafer, A., Sunkin, S. M., Oh, S. W., Bernard, A., … Ng, L. (2020). The Allen Mouse Brain Common Coordinate Framework: A 3D Reference Atlas. Cell, 181(4), 936-953.e20. 10.1016/J.CELL.2020.04.007

Wekselblatt, J. B., Flister, E. D., Piscopo, D. M., & Niell, C. M. (2016). Large-scale imaging of cortical dynamics during sensory perception and behavior. Journal of Neurophysiology, 115(6), 2852–2866. 10.1152/JN.01056.2015

Willeke, K. F., Fahey, P. G., Bashiri, M., Hansel, L., Blessing, C., Lurz, K.-K., Burg, M. F., Cadena, S. A., Ding, Z., Ponder, K., Muhammad, T., Patel, S. S., Deng, K., Guan, Y., Zhu, Y., Xiao, K., Han, X., Azeglio, S., Ferrari, U., … Albrecht, J. (2023). Retro-spective on the SENSORIUM 2022 competition. Proceedings of Machine Learning Research, 220, 314–333.

Yamins, D. L. K., Hong, H., Cadieu, C. F., Solomon, E. A., Seibert, D., & DiCarlo, J. J. (2014). Performance-optimized hierarchical models predict neural responses in higher visual cortex. Proceedings of the National Academy of Sciences of the United States of America, 111(23), 8619–8624. 10.1073/PNAS.1403112111/-/DCSUPPLEMENTAL/PNAS.201403112SI.PDF

Yatsenko, D., Fahey, Froudarakis, Reimer, Walker, Sinz, Cobos, & Tolias. (2018). trippy-monet/yatsenko-SfN2018-lowres.pdf at master · dimitri-yatsenko/trippy-monet. https://github.com/dimitri-yatsenko/trippy-monet/blob/master/yatsenko-SfN2018-lowres.pdf

Yoshida, T., & Ohki, K. (2020). Natural images are reliably represented by sparse and variable populations of neurons in visual cortex. Nature Communications 2020 11:1, 11(1), 1–19. 10.1038/s41467-020-14645-x

Zhang, Y., Rózsa, M., Liang, Y., Bushey, D., Wei, Z., Zheng, J., Reep, D., Joey Broussard, G., Tsang, A., Tsegaye, G., Narayan, S., Obara, C. J., Lim, J.-X., Patel, R., Zhang, R., Ahrens, M. B., Turner, G. C., S-H Wang, S., Korff, W. L., … Looger, L. L. (2023). Fast and sensitive GCaMP calcium indicators for imaging neural populations. 884 | Nature |, 615. 10.1038/s41586-023-05828-9

Zhuang, J., Ng, L., Williams, D., Valley, M., Li, Y., Garrett, M., & Waters, J. (2017). An extended retinotopic map of mouse cortex. ELife, 6, 1–29. 10.7554/eLife.18372

