## Supplementary figures and images for "Mapping the visual cortex with Zebra noise and wavelets"

### Supplementary Video 1

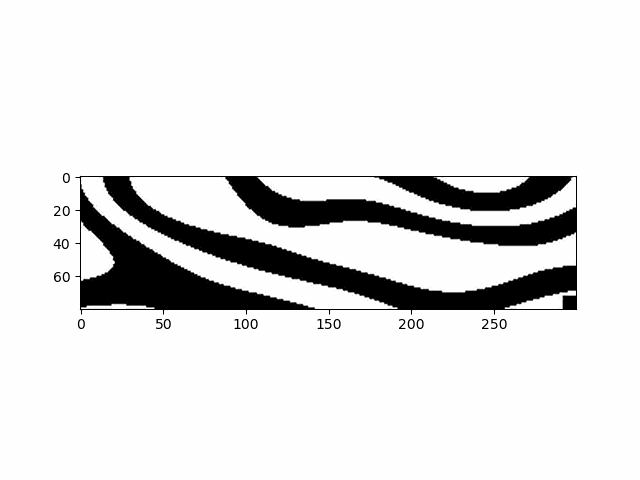
